# Native nucleosome-positioning elements for the investigation of nucleosome repositioning

**DOI:** 10.1101/2025.01.17.633597

**Authors:** Ruo-Wen Chen, Shane D. Stoeber, Ilana M. Nodelman, Hengye Chen, Lloyd Yang, Gregory D. Bowman, Lu Bai, Michael G. Poirier

**Affiliations:** Ohio State Biochemistry Graduate Program, The Ohio State University, Columbus, OH 43210, USA; Department of Biochemistry and Molecular Biology, Center for Eukaryotic Gene Regulation, The Pennsylvania State University, University Park, PA 16802, USA; TC Jenkins Department of Biophysics, Johns Hopkins University, Baltimore, MD 21218, USA; Department of Physics, The Ohio State University, Columbus, OH 43210, USA; Department of Physics, The Pennsylvania State University, University Park, PA 16802, USA; Department of Chemistry & Biochemistry, The Ohio State University, Columbus, OH 43210, USA

**Keywords:** *nucleosome positioning*, *chromatin biology*, *transcription factors*, *ATP-dependent chromatin remodeling*, *nucleosome dynamics*

## Abstract

Nucleosome repositioning is essential for establishing nucleosome-depleted regions (NDRs) to initiate transcription. This process has been extensively studied using structural, biochemical, and single-molecule approaches, which require homogenously positioned nucleosomes. This is often achieved using the Widom 601 sequence, a highly efficient nucleosome positioning element (NPE) selected for its unusually strong binding to the H3-H4 histone tetramer. Due to the artificial nature of 601, native NPEs are needed to explore the role of DNA sequence in nucleosome repositioning. Here, we characterize the position distributions and nucleosome formation free energy for a set of yeast native nucleosomes (YNNs) from *Saccharomyces cerevisiae*. We show these native NPEs can be used in biochemical studies of nucleosome repositioning by transcription factors (TFs) and the chromatin remodeler Chd1. TFs could directly reposition a fraction of nucleosomes containing native NPEs, but not 601-containing nucleosomes. In contrast, partial unwrapping was similar for 601 and native NPE sequences, and the rate of ATP-dependent remodeling by Chd1 was within the range of the fast and slow directions of the 601 nucleosomes. This set of native NPEs provides an alternative to the 601 NPE that can be used for probing the repositioning of nucleosomes that contain native DNA sequences.

## INTRODUCTION

All eukaryotic genomes are organized into long repeats of the highly conserved DNA-protein complex, the nucleosome, to form chromatin fibers. Each nucleosome contains ∼147 bp of genomic DNA wrapped 1.65 times around a histone protein octamer (1). Nucleosomes regulate gene transcription by controlling DNA accessibility to the transcription machinery (2, 3). In particular, the DNA positioning of nucleosomes relative to transcription factor binding sites is a key mechanism for the regulation of a gene’s transcriptional state. Often, nucleosome positioning creates nucleosome-depleted regions (NDRs) within gene promoters and enhancers, which increases the accessibility to help initiate transcription (4, 5). Studies have shown that NDR formation is often initiated by pioneer transcription factors (TFs), yet nucleosome positioning can also be driven by canonical TFs (6, 7). TFs then function with transcription co-activators, including histone acetyltransferases and ATP-dependent nucleosome remodelers, to open chromatin for the recruitment of the transcription machinery (8–11). While the generation of NDRs within gene promoters and enhancers is well established (12–14), how particular nucleosomes are directionally shifted remains enigmatic.

Nucleosome positions are influenced not only by TFs and co-activators but also by the underlying DNA sequence (15). This sequence dependence has been exploited to produce well-positioned nucleosome samples for biochemical and biophysical studies, like the NPE within the *Lytechinus variegatus* 5S ribosomal RNA (5SR) gene (16). In a foundational study exploring how DNA sequence affects nucleosome positioning, Lowary and Widom identified artificial DNA sequences with high histone octamer binding affinities that position nucleosomes with remarkable efficiency (17). Among these, the “601” sequence exhibits the strongest histone octamer binding and has become the standard for preparing homogenously positioned nucleosomes. The 601 sequence’s strong positioning capability has facilitated numerous studies on nucleosome structure and dynamics, including investigations of TF binding to partially unwrapped nucleosomes (18, 19) and ATP-dependent nucleosome remodeling (20–22).

Recent single-molecule measurements that quantified the rate of ATP-independent nucleosome repositioning revealed that repositioning of 601 nucleosomes is significantly more limited than nucleosomes that contain the DNA sequence from the mouse *Lhb* gene promoter (23). Additionally, canonical and pioneer TF binding within partially unwrapped 601 nucleosomes do not lead to changes in the dyad positioning (18, 24), while pioneer TF binding within a native NPE from the CX3CR1 gene results in nucleosome repositioning (25). Furthermore, studies of the ATP-dependent remodeler Chd1 found that inserting poly-A tracts and changing orientations of 601 nucleosomes resulted in very different remodeling rates (20, 26), revealing that DNA sequence can influence nucleosome remodeling. In combination, these findings underscore the need for a broader library of well-characterized native NPEs for investigating nucleosome repositioning and dynamics in physiologically relevant contexts.

Here, we establish a set of NPEs derived from the *Saccharomyces cerevisiae* genome for nucleosome repositioning studies, which are referred to as Yeast Native Nucleosome (YNN) sequences 1 to 7. We characterized the nucleosome position distribution and the nucleosome formation free energy for each YNN sequence. Interestingly, the homogeneity of nucleosome positioning does not correlate with the free energy. In fact, all YNNs have similar nucleosome formation free energies that were each significantly less stable than the 601 sequence. We then used a subset of these NPEs (YNN1–YNN4) to investigate nucleosome repositioning. Nucleosomes containing native NPEs are partially repositioned by TF binding, while 601 nucleosomes are not. In contrast, TF binding equilibrium within partially unwrapped nucleosomes was similar between 601 and YNN1. In addition, while the NPE also influenced the ATP-dependent nucleosome repositioning by Chd1, the remodeling efficiencies of YNN1 – YNN4 nucleosomes are largely similar to 601 nucleosomes. This study provides a set of native NPEs that will be valuable for nucleosome repositioning studies. Furthermore, these results indicate that it is important to consider native NPEs for ATP-independent nucleosome repositioning, that the NPE can quantitatively impact nucleosome partial unwrapping, and provide evidence that native NPE can affect the rate and directionality of nucleosome sliding by chromatin remodelers.

## METHODS

### Identification of NPE candidates

NPE candidate selection was done by analyzing and comparing published data of *in vivo* budding yeast nucleosome positions and position data for nucleosomes prepared by *in vitro* salt dialysis reconstitutions (27, 28). We generated a 147 bp nucleosome profile where the middle 80 bp is fully protected, while the entry and exit regions exhibit partial protection (**Figure S1A**). To identify nucleosome-like particles, we computed the correlation between this profile and 147 bp sliding windows across the genome-wide nucleosome occupancy data both *in vivo* and *in vitro*, using a 5 bp step size (example shown in **Figure S1B**). Potential nucleosome dyad positions were identified as peaks in the correlation curves, with the following criteria: 1) the peak amplitude must be greater than 0, and 2) the nucleosome occupancy at the peak position must exceed 1, which is the genome-wide average.

After collecting all the correlation values and dyad positions *in vivo* and *in vitro*, we compared them to find nucleosomes that are aligned between these two datasets. Specifically, the aligned nucleosomes are required to have a correlation >0.7 both *in vivo* and *in vitro*, and the distance between their dyad positions ≤ 20 bp. Using these criteria, we identified 6,675 aligned nucleosomes across the genome. **Figure S1C** shows the distribution of the scores of the aligned nucleosomes, calculated as the sum of their correlation values *in vivo* and *in vitro*. Seven candidate sequences, YNN1 to YNN7, were selected (**Table S1**), which have typical scores among this group that span a wide range (**Figure S1C**). The correlations and dyad distances of YNN1-7 are listed in **Table S2**.

### Reb1 preparation

Reb1 was generated and purified as previously mentioned (29). Reb1 was cloned into pHIS8 plasmid and over-expressed in *Escherichia coli* BL21(DE3) strain when OD_600_ reached 0.4-0.6 through IPTG induction for 3 hours under 37 ⁰C. Cells were then collected by centrifugation (Beckman Coulter Avanti) at 4000xg for 10 minutes and resuspended in lysis buffer (50 mM NaH_2_PO_4_ pH 8.0, 300 mM NaCl, 10 mM imidazole, 1 mM PMSF, 1 mM DTT) for cell lysis through sonication. Supernatant from the cell lysate was collected by another round of centrifugation at 4000xg for 10 minutes and loaded onto a 5 ml HisTrap HP column (Cytiva). Using fast protein liquid chromatography (abbreviated as FPLC, ÄKTA Pure from Cytiva), we eluted the protein with elution buffer (50 mM NaH_2_PO_4_ pH 8.0, 300 mM NaCl, 250 mM imidazole, 1 mM PMSF, 1 mM DTT). The eluted sample was further purified over a size-exclusion column (Superdex 200 from Cytiva) pre-equilibrated in Reb1 storage buffer (20 mM HEPES pH 7.5, 350 mM NaCl, 1% Tween-20). Reb1 was aliquoted with additional glycerol (10% final concentration), flash frozen in liquid nitrogen, and then stored at −80 ⁰C. The purified Reb1 was examined by sodium dodecyl-sulfate polyacrylamide gel electrophorsis (SDS-PAGE) with 8% acrylamide and stained with Coomassie brilliant blue (**Figure S2B**).

### Cbf1 preparation

Cbf1 was generated and purified as previously reported (30). Cbf1 was cloned in pET28a plasmid and expressed in *Escherichia coli* BL21(DE3) strain through 0.5 mM IPTG induction for 4 hours at 37 ⁰C when OD600 reached 0.4-0.6. Cells were collected and resuspended in lysis buffer (50 mM Na_2_HPO_4_ pH7.5, 300 mM NaCl, 5 mM imidazole, 10% glycerol, 1 mM PMSF, 20 µg/ml pepstatin, 20 µg/ml leupeptin) for sonication. The supernatant was separated from the cell debris through centrifugation (Beckman Coulter Avanti) at 23000xg at 4 ⁰C for 20 minutes and manually loaded onto a 5 ml HisTrap HP column (Cytiva). The column was sequentially flushed with 40 ml lysis buffer, 120mL wash buffer (50 mM Na_2_HPO_4_ pH 7.5, 300 mM NaCl, 60 mM imidazole, 10 % glycerol, 1 mM PMSF, 20 µg/mL pepstatin, 20 µg/ml leupeptin), and elution buffer (50 mM Na_2_HPO_4_ pH 7.5, 300 mM NaCl, 340 mM imidazole, 10 % glycerol, 1 mM PMSF, 20 µg/ml pepstatin, 20 µg/mL leupeptin). The eluted Cbf1 was buffer-exchanged into storage buffer (50 mM Na_2_HPO_4_ pH 7.5, 300 mM NaCl, 10 % glycerol, 1 mM PMSF, 20 µg/ml pepstatin, 20 µg/ml leupeptin) through Amicon Ultra Centrifugal Filters (Millipore Sigma), flash frozen in liquid nitrogen, and stored in −80 ⁰C. Purified Cbf1 was examined on the SDS-PAGE via Coomassie blue staining (**Figure S2C**).

### Pho4 preparation

Pho4 was prepared and purified as previously mentioned (30). Pho4 was over-expressed in *Escherichia coli* BL21(DE3) strain through 1 mM IPTG induction when OD600 reach 0.4-0.6 and harvested after incubation for 3 hours under 37 ⁰C. Cells were collected and resuspended in lysis buffer (20 mM Tris-HCl pH 8.0, 0.5 M NaCl, 10 mM imidazole, 0.2% Triton X-100, 5 mM DTT, 1 mM PMSF, 20 µg/mL pepstatin, 20 µg/mL leupeptin, 2.1 mM benzamidine). Cells were lysed through sonication, and the supernatant was separated from the cell lysates through centrifugation (Beckman Coulter Avanti) at 23000xg for 20 minutes at 4 ⁰C and loaded onto a 5 mL HisTrap HP Ni-NTA column (Cytiva). The HisTrap column was washed with 150 mL of wash buffer A with detergent (20 mM Tris-HCl pH 8.0, 0.5 M NaCl, 5 mM imidazole, 0.02% Triton X-100, 5 mM DTT), followed by 150 mL of wash buffer B without detergent (20 mM Tris-HCl pH 8.0, 0.5 M NaCl, 5 mM imidazole, 5 mM DTT), and gradually eluted with elution buffer (20 mM Tris-HCl pH 8.0, 0.5 M NaCl, 500 mM imidazole). The eluted protein was further purified through a size exclusion column (Superdex 200 from Cytiva) into the storage buffer (40 mM HEPES-NaOH pH7.4, 200 mM potassium acetate, 1 mM EDTA). Glycerol was added to the purified Pho4 to 10% final concentration. The final product was examined on SDS-PAGE and stained with Coomassie (**Figure S2D**).

### DNA preparation

All DNA constructs were cloned in pUC19 and amplified through polymerase chain reaction (PCR). The Cy3-labeled DNA was either purchased as 5’-labeled oligonucleotides from Integrated DNA Technologies (IDT) or internally labeled through the Cy3-NHS ester-activated crosslinking to amino-modifier C6 dT (**Table S3**). The internal-labeled oligonucleotides were purified through a reverse phase column (C18 from Thermo Fisher Scientific) from free Cy3. PCR was carried out using *Pfu* DNA polymerase, which contains an exonuclease proofreading activity. Amplified DNA was purified through an anion exchange chromatography column (Mono Q from Cytiva) using high-performance liquid chromatography (abbreviated as HPLC, Agilent). Purified DNA was buffer-exchanged into 0.5x TE buffer (5 mM Tris-HCl pH8.0, 0.5 mM EDTA) by Amicon Ultra Centrifugal Filters (Merck Millipore).

### Histone octamer preparation

Human histones (H2A with K119C, H2B, H3(C110A) and H4) were obtained from The Histone Source. Histone octamers were refolded with a 1.3:1 molecule ratio of H2A and H2B to H3 and H4 through dialyzing from the unfolding buffer (7M guanidine hydrochloride, 20 mM Tris-HCl pH 7.5, 10 mM DTT) to the refolding buffer (2M NaCl, 10 mM Tris-HCl pH 7.5, 1 mM EDTA pH8.0, 5 mM 2-Mercaptoethanol). The Cy5-labeled histone octamers were generated through the maleimide-thiol reaction at H2AK119C, and the reaction was quenched by adding 10 mM DTT. The refolded histone octamers were purified by FPLC gel-filtration chromatography (Superdex 200 from Cytiva). Final products were examined by 16% SDS-PAGE and stained with Coomassie brilliant blue.

*Xenopus laevis* histones (H2A, H2B(S53C), H3(C110A/M120C), H4) were expressed in *E. coli* and purified from inclusion bodies as previously described (31). Purified histones were unfolded at room temperature in an unfolding buffer and, after one hour, mixed in a 1:1:1:1 molar ratio. The histone mixture was dialyzed at 4°C against three changes of refolding buffer, and, the next day, concentrated and purified over a HiLoad Sephadex S200 16/600 (Cytiva). Fractions were assessed by SDS-PAGE (18%), and those containing octamer were pooled and concentrated. Following the addition of 10% glycerol (Cf), the sample was aliquoted, flash-frozen in liquid nitrogen, and stored at −80°C. This octamer was used for nucleosomes employed in site-directed cleavage and nucleosome mapping studies (**Figure 5 and S10**).

### Recombinant nucleosome reconstitution

Histone octamers and DNA segments were reconstituted into nucleosomes with a molar ratio of 1:2 through double salt dialysis from the high salt buffer (2M NaCl, 5 mM Tris-HCL pH8.0, 0.5 mM EDTA, 1 mM benzylamine) to the refolding buffer (5 mM Tris-HCL pH8.0, 0.5 mM EDTA, 1 mM benzylamine) as previously described (12). The reconstituted nucleosomes are purified through sucrose gradient centrifugation with an ultracentrifuge (Beckman Coulter) spun at 41000 rpm for 22 hours under 4 ⁰C. The purification used a 5% to 30% sucrose gradient in 0.5x TE buffer that was prepared with a Gradient Master (Biocomp). The fractions of sucrose gradient were collected via a Model 2110 Fraction Collector (Bio-Rad) and examined on a 5% native polyacrylamide gel at 300 V for 1 hour at room temperature (**Figure S3**). Purified nucleosomes were buffer-exchanged into the storing buffer (5 mM Tris-HCl pH8.0, 0.5 mM EDTA, 20% glycerol), flash frozen with liquid nitrogen, and stored in −80 ⁰C.

For Figure 5 and Supplemental Figures 10 and 11, nucleosomes were prepared as described (32, 33). Specifically, double-labeled DNA, Cy5-40-YNN1-40-FAM, was generated from pUC19-YNN1 template in 18 ml of PCR and purified over a Miniprep cell apparatus (Bio-Rad). This DNA was combined in a 1:1 ratio with purified *Xenopus laevis* histone octamer (H2Awt, H2B-S53C, H3-M120C, H4wt) and reconstituted using salt gradient dialysis. Nucleosomes were purified over a 7% (60:1) acrylamide column using a Prep Cell (Bio-Rad) and eluted in 10 mM Tris-HCl, pH 7.5, 2.5 mM KCl, 1 mM EDTA (pH 8.0), and 1 mM DTT. Collected fractions were run on 7% (60:1) native gels and bands visualized on the Typhoon 5 scanner. Selected fractions were pooled, concentrated, and 100% glycerol was added to a final concentration of 20%. Nucleosomes were aliquoted and flash frozen in liquid nitrogen and stored at −80°C.

### Competitive nucleosome reconstitution

Competitive nucleosome reconstitution was carried out as previously described (34). Nucleosomes were reconstituted with 3 ug unlabeled histone octamer, 200 ng Cy3-labeled sample DNA, and 20 ug unlabeled buffer DNA through the double salt dialysis method. The buffer DNA was a 147bp DNA segment from the ampicillin resistance gene (ampR) in the pUC19 plasmid (**Table S3**). The results of reconstitution were analyzed by native polyacrylamide gel electrophoresis (PAGE) with 5% polyacrylamide gel that was imaged with the Typhoon biomolecular imager (Cytiva) at the Cy3 channel.

### Electrophoresis mobility shifting assay (EMSA)

EMSA was performed with a 5% native polyacrylamide gel in 0.3x TBE (30 mM Tris base, 27 mM boric acid, and 0.3 mM EDTA) buffer. Native gels were pre-run at 4⁰C with 300 V for 90 minutes and then run for another 90 minutes under the same conditions after loading the samples. A titrated amount of Reb1 was incubated with 0.5 mM nucleosomes for 20 minutes in T130 buffer (10 mM Tris-HCl pH 8.0 with 130 mM NaCl, 10% glycerol, and 0.0075% Tween-20) at room temperature before running the gel. The gels were imaged with a Typhoon biomolecular imager. EMSA gels were quantitatively analyzed through ImageJ (NIH) software. The statistical analysis was done with the GraphPad Prism software. The binding measurements were fit to the Hill equation to obtain S_1/2_.

### Sample preparation of MNase-seq

Nucleosome binding experiments were performed by incubating 60 nM nucleosome with 0 nM or 60 nM of TFs (Reb1, Cbf1, or Pho4) for 20 minutes at room temperature. Nucleosomes-TF binding was confirmed on a 5% acrylamide native gel. The incubated samples were then digested with 0.3 U, 1 U, 3 U, 10 U of MNase (New England Biolabs) for 8 minutes at 37 ⁰C and immediately quenched by adding quench buffer (0.1 % SDS, 20 mM EDTA pH8.0, 0.01 U proteinase K). Proteinase K digestion was carried out by incubating the quenched samples for 1 hour at 50 ⁰C. The final digested samples were analyzed by agarose gel electrophoresis with a 1.5% gel in 0.5x TAE buffer (20 mM Tris base, 10 mM acetic acid, 0.5 mM EDTA pH 8.0) (**Figure S4**). The digest DNA samples were then purified by agarose gel electrophoresis and then gel extraction with a Freeze ‘N Squeeze DNA extraction kit (Bio-Rad).

### NGS library preparation and MNase-seq

After gel purification of the MNase digested samples, sequencing libraries were prepared using NEBNext ® Ultra™ II DNA Library Prep Kit for Illumina® (Catalog # E7645L) according to the manufacturer’s instructions. The sequencing libraries were sequenced on a NextSeq 2000 machine in 2 x 50 bp paired-end sequencing mode at the Genomics Research Incubator at The Pennsylvania State University. Nucleosome occupancy plots were created by aligning the sequencing reads to the corresponding reference sequence (Widom 601 or YNN with or without factor binding site) with Bowtie2 on galaxy in paired-end mode. Unmapped and unpaired reads were removed using Samtools sort. Next, nucleosome coverage was calculated from the read counts normalized to total reads in BPM using bamCoverage. The minimum and maximum fragment length was set to 130 bp and 160 bp respectively, unless otherwise stated to enrich reads for mono-nucleosome fragments. The dyad position for each sequence was calculated using bamCoverage –MNase in which only the three center bp of each fragment was used for analysis and were normalized to total reads in BPM. Bigwig files were converted to BedGraph format for further analysis.

### Förster resonance energy transfer (FRET)

Nucleosome wrapping and shifting events were observed by monitoring the efficiency of energy transfer between the donor (Cy3) and the acceptor (Cy5) that were labeled within the nucleosome. While Cy5 was labeled at H2AK119C, Cy3 was externally labeled at the 5’ end or internally labeled within the linker DNA. The binding experiments were performed by incubating 0.5 nM nucleosomes with a titrated amount of TF for 20 minutes at room temperature and immediately measured the fluorescence spectra using FluoroMax 4 fluorimeter (Horiba). Cy3 was excited at 510 nm and Cy5 was excited at 610 nm, while their fluorescence emission was measured from 530 to 750 nm and from 630 nm to 670 nm, respectively. The ratio_A_ method (35) was used to calculate the FRET efficiency from the obtained spectra as previously described (19). All the FRET measurements were replicated three times, and the error bars represent the standard error from the replicates. The binding measurement between Reb1 and proximally labeled nucleosomes were fit to the Hill equation to obtain the S_1/2_.

### Chd1 preparation

*Saccharomyces cerevisiae* Chd1 (amino acids 118-1274) was prepared as previously described (31). Briefly, Chd1 was expressed overnight at 18°C following IPTG induction in BL21-DE3 RIL cells. Cells were lysed by sonication and spun. The supernatant was then passed over a HisTrap HP (Cytiva) column equilibrated in 50 mM Tris-HCl, pH 7.5, 500 mM NaCl, 10 mM imidazole, and 10% glycerol. Chd1 was eluted in the equilibration buffer with 175 mM imidazole. Pooled fractions were diluted five-fold to reduce NaCl concentration to 100 mM, spun, and loaded onto a cation exchange (HiTrap SPFF, Cytiva) column. Chd1 was eluted via a linear salt gradient of 100-500 mM NaCl (in 30 mM Tris-HCl, pH 7.5 with 10% glycerol). Chd1 was digested overnight by Precision Protease and then repassed over the nickel column, concentrated, and further purified over a HiLoad Sephadex S200 16/600 (Cytiva) equilibrated in 30 mM Tris-HCl, pH 7.5, 150 mM NaCl, and 10% glycerol. The final protein pool was concentrated, aliquoted, and flash-frozen in liquid nitrogen.

### Site-directed DNA cleavage and histone mapping

Cy5-40-YNN1-40-FAM nucleosomes with octamer containing *Xenopus laevis* H3M120C, H2A, H2B-S53C, and H4 were incubated in the dark with 250 µM 4-azidophenacyl bromide (APB) for 2.5-3 hours at room temperature as previously described (21, 36). The labeling reaction was quenched with 5 mM DTT, and a multi-reaction mix was prepared with 150 nM budding yeast Chd1 (amino acids 118-1274) and 150 nM APB-labeled nucleosomes in mapping reaction buffer: 20 mM Tris-HCl, pH 7.5, 50 mM KCl, 5 mM MgCl2, 100 ng/µl BSA, 5 mM DTT, and 10% glycerol. A minus ATP control reaction was removed from the mix, and then 2 mM ATP was added and 50 µl time points were quenched with 100 µl of mapping reaction buffer (-MgCl2) plus 25 mM EDTA, pH 8.0 and placed on ice. Samples were then UV (302 nm) irradiated for 1 minute and 15 seconds.

For Chd1 mapping experiments, single cysteine ATPase mutants N459C (Lobe 1) or V721C (Lobe 2) were incubated at room temperature for 2.5-3 hours in the dark with 400 µM APB as described in (31). A three-fold excess of APB-labeled Chd1 (450 nM final concentration) was added to 150 nM Cy5-40-YNN1-40-FAM nucleosomes and incubated for 12 minutes in the presence of either AMPPNP or ADP-BeF3 in mapping reaction buffer. Samples were then UV irradiated.

Following UV irradiation, Chd1 and histone crosslinking samples were treated equivalently in the manner of (36). Specifically, 150 µl of 20 mM Tris-HCl, pH 8.0, 50 mM NaCl, and 0.2% SDS buffer was added, samples were mixed, and incubated at 70°C for 20 min. Next, 300 µl of 5:1 phenol:chloroform (cat# P1944 Sigma) was added to each sample, vortexed, spun, and ∼200 µl of the top layer was removed and discarded. Following this, 300 µl of a mixture of 1 M Tris-HCl, pH 8.0/1% SDS was added to each sample, vortexed, and spun, and again, the top ∼200 µl was discarded. This step was repeated for a total of 4 times. Samples were precipitated overnight on ice after the addition of 1 µl of 10 mg/ml glycogen, 35 µl 3 M NaOAc, pH 5.3, 750 µl 100% EtOH. The next morning, samples were spun (4°C) for 30 mins, 15000 rpm, and washed twice with 70% EtOH. Samples were dried in the hood for a minimum of 1 hour, then pellets were dissolved in 100 µl 20 mM ammonium acetate, 2% SDS, 0.1 mM EDTA, pH 8.0, thoroughly vortexed, and spun for 10 mins. The supernatant was transferred to fresh tubes, and heated at 90°C for 10 mins, then 0.1 N NaOH concentration final was added and samples were incubated another 40 mins at 90°C. Following this, 105 µl 20 mM Tris-HCl, pH 8.0, 1.0 µl 2 M MgCl2, 4.5 µl 2N HCl, and 480 µl 100% EtOH was added to precipitate DNA. Samples were left overnight at −20°C, and the next morning, samples were spun, washed two times in 70% EtOH, and allowed to dry. Pellets were resuspended in 4 µl formamide loading buffer and loaded onto an 8% (19:1) acrylamide/8M urea denaturing gel. Gel was run at 65 Watts for up to ∼1 hour 40 mins and scanned on a Typhoon 5 scanner (Cytiva). A sequencing ladder was generated using Therminator (NEB) and a mixture of standard nucleotides spiked with individual acyclonucleotides (NEB).

### Chd1 remodeling assay

Nucleosome sliding reactions by Chd1 were analyzed as previously reported (20). Before initiating remodeling reactions, 10 nM nucleosomes were pre-incubated on ice with Chd1 in 1x sliding buffer (20 mM Tris-HCl, 50 mM KCl, 5 mM MgCl_2_, 5% sucrose, 1mM DTT, 0.1 mg/mL BSA) for 10 minutes in a total 10 µL volume. For time-course experiments, 10 nM nucleosomes were incubated with 10 nM Chd1 in the remodeling reaction. As for the other experiments, 10 nM nucleosomes were titrated with an increasing concentration of Chd1 (0 nM, 1 nM, 3 nM, 10 nM, 30 nM). Sliding reactions were initiated by adding 1 mM ATP and incubated at room temperature for 0, 1, 2, 4, 8, 16, 32 minutes for kinetics studies. Each reaction was then quenched by 60 µL 1x quench buffer (20 mM Tris-HCl, 50 mM KCl, 5 mM MgCl_2_, 5% sucrose, 1mM DTT, 0.1 mg/mL BSA, 25 mM EDTA, 9 µg salmon sperm DNA) and analyzed by EMSAs with 5% polyacrylamide native gels. Gels were scanned with a Typhoon biomolecular imager, and the fraction of each shifted band was quantified with ImageJ (NIH) software. GraphPad Prism software was used to generate plots and carry out statistical analysis. All time course measurements of Chd1 remodeling were done in triplicate and were fit to the function: fraction shifted = F_i_ + (F_f_ – F_i_)*(1-exp(-k*t)), where t represents time, F_i_ is the initial shifted faction, F_f_ is the final fraction shifted, and k is the rate constant (Figure 7C). The error bars (Figure 7) are the standard error of the three repeats.

## RESULTS

### Identifying candidate nucleosome-positioning elements (NPEs) from the native yeast genome

To identify natural sequences that form well-positioned *in vitro* reconstituted nucleosomes, we utilized previously published *in vitro* and *in vivo* nucleosome occupancy data over the yeast genome measured by micrococcal nuclease digestion and high throughput sequencing (MNase-seq) (27, 28). The *in vitro* data was generated from nucleosomes assembled by salt gradient on yeast genomic DNA using a 0.4:1 histone-to-DNA mass ratio, significantly lower than the 1:1 mass ratio in typical eukaryotic cells. Partly due to the lower nucleosome density in this dataset, most nucleosome positions appeared fuzzy. We reasoned that the sequences associated with well-positioned nucleosomes under these conditions would be strong candidates for NPEs. To ensure physiological relevance, we refined our selection by filtering for sequences that also form well-positioned nucleosomes *in vivo*. In brief, we correlated the genome-wide *in vivo* and *in vitro* nucleosome occupancies with a “standard” nucleosome occupancy curve to identify nucleosome-like particles and their dyad positions (see Methods for details). Using the criteria that the *in vivo* and *in vitro* correlations both exceed 0.7 and that the dyad positions are within 20bp, we identified 6675 potential NPEs across the yeast genome. From these, we selected seven NPE candidates spanning a range of correlation and dyad alignment values, which we named Yeast Native Nucleosome 1-7 (YNN 1-7) (**Figure 1, S1, Table S1**). Despite being selected solely based on nucleosome occupancy, all of these sequences are located within open reading frames, likely because yeast intergenic regions tend to contain poly(A/T) stretches that have low intrinsic histone affinities (37, 38).

**Figure 1.**
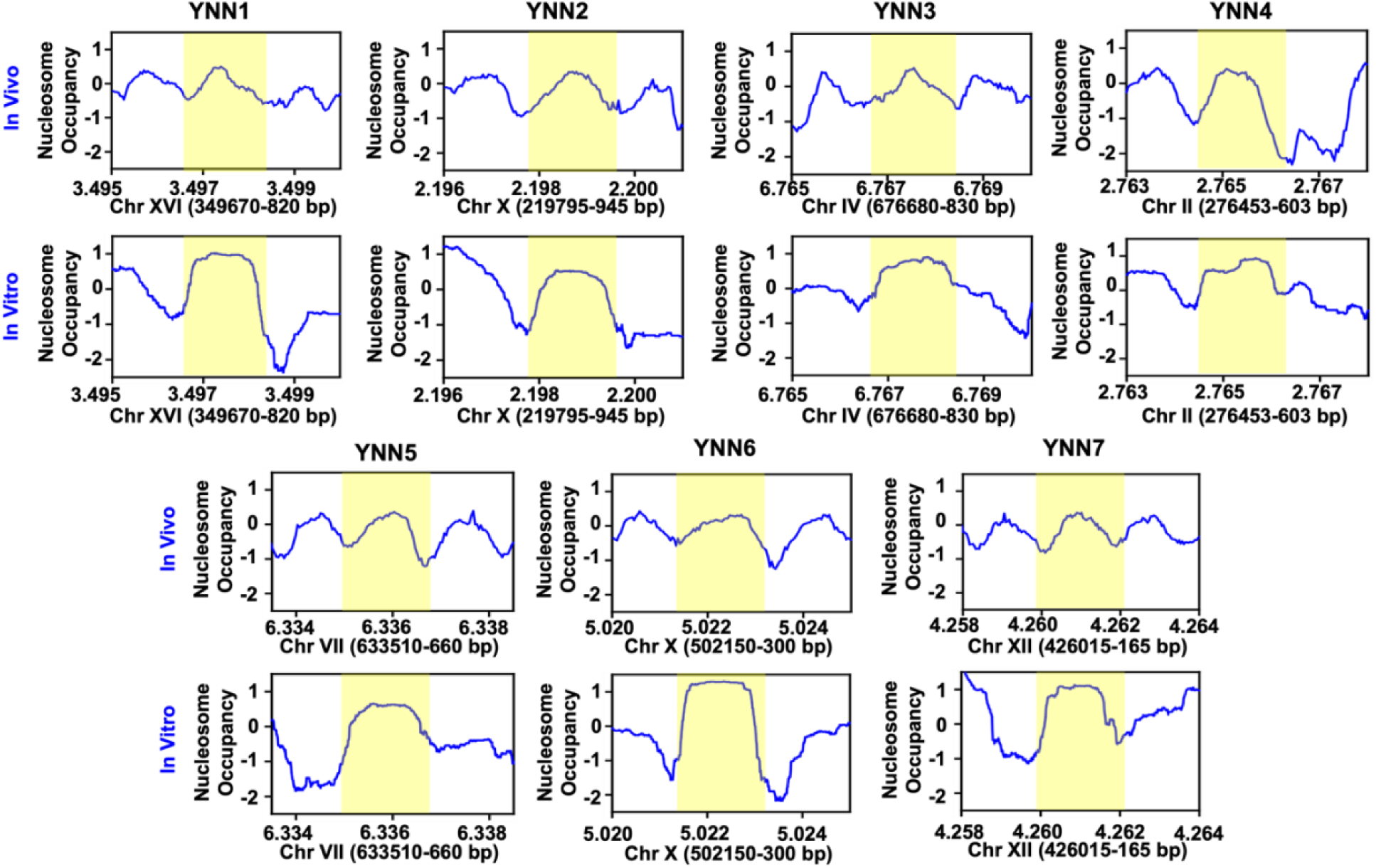
Identifying candidate nucleosome-positioning elements (NPEs) from the budding yeast genome. Nucleosome occupancy measured *in vivo* and *in vitro* through global MNase-seq in budding yeast genome. The yellow regions are the 147 bp YNN sequences.

### A subset of NPE candidates form well-positioned nucleosomes by salt dialysis reconstitutions

After identifying these seven native yeast DNA sequences, we investigated how well these sequences position nucleosomes following *in vitro nucleosome* reconstitution. To investigate this, we prepared 247 base pair (bp) long DNA molecules that contain a budding yeast genomic sequence that is centered according to the estimated nucleosome position *in vivo* (**Figure 2A**). We then separately reconstituted nucleosomes via salt dialysis with each YNN DNA molecule and Cy5-labeled histone octamer (**Figure 2B and S3**). The reconstituted nucleosomes were then analyzed by electromobility shift assays (EMSAs), which are sensitive to nucleosome position where the electromobility is slower as the nucleosome is more centered (**Figure 2C**).

**Figure 2.**
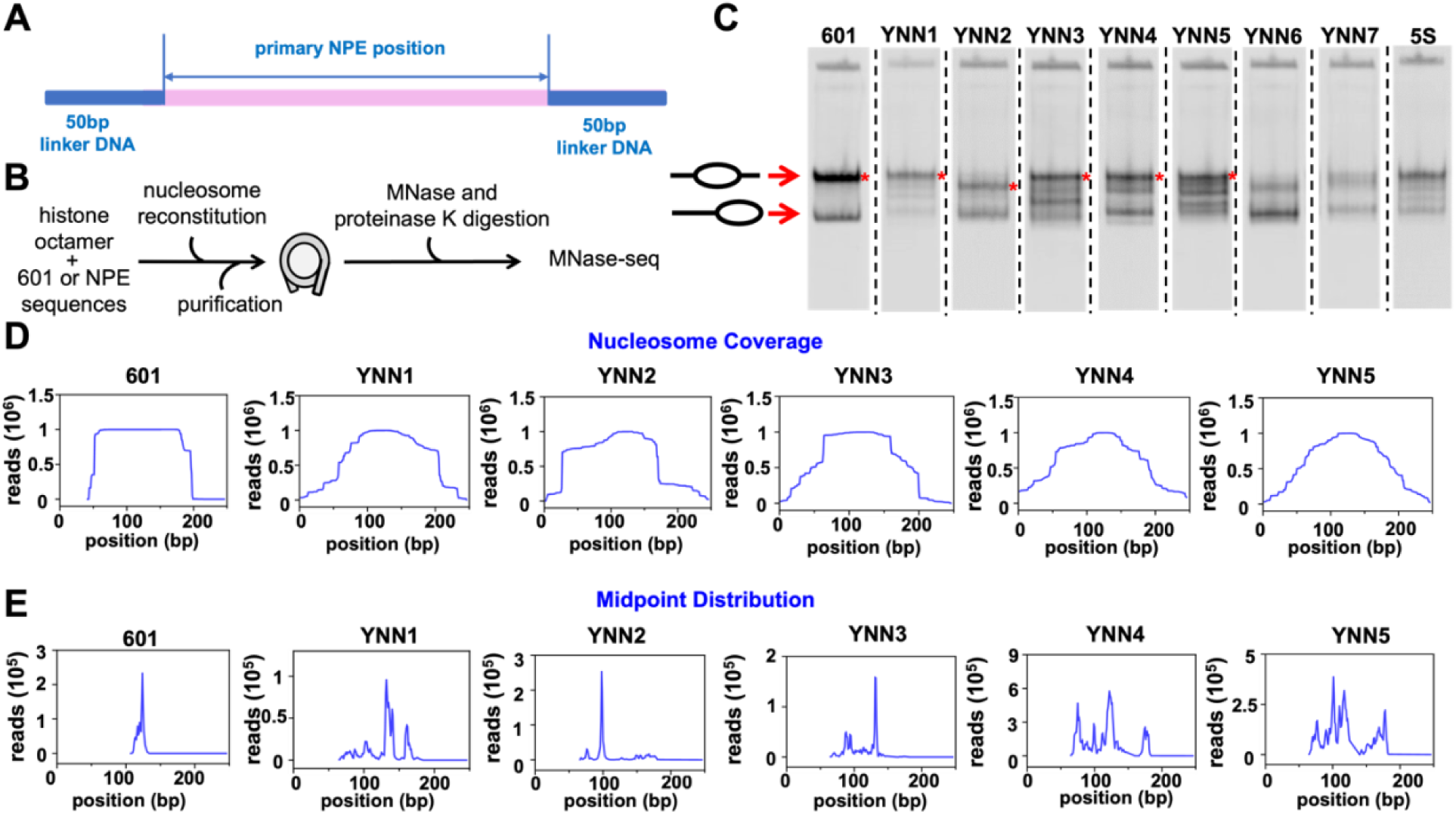
A subset of NPE candidates form well-positioned nucleosomes by salt dialysis reconstitutions. **(A)** 247bp DNA construct used for generating recombinant mono-nucleosomes. Each DNA contains 50bp linker DNA at both ends of the 147bp primary NPE position. **(B)** Workflow of nucleosome reconstitution and MNase-seq. Nucleosomes were generated through salt dialysis from 2M NaCl to 0M NaCl, which was followed by sucrose gradient centrifugation to purify nucleosomes from excess DNA. Cy5-labeled histone octamers were used for detecting the nucleosome positioning in the EMSAs. **(C)** Cy5 fluorescence image of an EMSAs for quantifying nucleosome positioning of different NPEs following salt dialysis reconstitution. The different mobility bands are indicated as center-positioned nucleosomes (upper) and end-positioned nucleosomes (lower). The red asterisks denote center-position nucleosomes. The images of each lane are from the same gel and were rearranged to be in order of the naming convention. **(D** and **E)** Nucleosome coverage and midpoint distributions of nucleosomes containing 601 and YNN1-5, which were determined from the MNase-seq measurements.

We included the Widom 601 (17) and *Lytechinus variegatus* 5S (16) DNA sequences as standards for nucleosome positioning and as references for both center (slowest electrophoretic mobility) and end (highest electrophoretic mobility) positioned nucleosomes. The 601 sequence was originally selected for its high propensity to form nucleosomes by salt dialysis reconstitutions, which resulted in a sequence that forms highly stable and positioned nucleosomes (17). *Lytechinus variegatus* 5S DNA is a native sequence that was identified to form well-positioned nucleosomes by salt dialysis and has been used extensively for *in vitro* measurements (16). Following salt dialysis reconstitution, the DNA molecule with the 601 sequence forms a single slow electrophoretic mobility band, which contains nucleosomes positioned at the center of the 601 sequence (**Figure 2C**). In addition, a minor high mobility band is also observed, which likely contains nucleosomes that form at the end of the 247 bp DNA molecule (39). The DNA molecule with the 5S sequence forms a dominant band with slow electrophoretic mobility similar to the 601 center-positioned and end-positioned nucleosomes (**Figure 2C**). However, there are additional bands with intermediate electrophoretic mobilities that indicate there are multiple nucleosome positions that are formed with DNA that contain this 5S sequence.

Interestingly, the YNN sequences vary significantly in the fraction of center-positioned nucleosomes and the number of bands with intermediate gel mobilities (**Figure 2C**). YNN1 forms center-positioned nucleosomes with a small fraction of off-center nucleosome positions, similar to nucleosomes reconstituted with 5S DNA. YNN2 forms a predominant nucleosome position that is off-centered and a second species that is likely end-positioned nucleosomes. YNN3, YNN4, and YNN5 form a significant fraction of center-positioned nucleosomes accompanied by multiple additional positions. Two of the identified sequences did not efficiently form center position nucleosomes. YNN6 largely forms end-positioned nucleosomes with a minor fraction of off-center positioned nucleosomes, while YNN7 did not efficiently form reconstituted nucleosomes. These results indicate that YNN1-5 could be used to reconstitute nucleosomes with different distributions of positioned nucleosomes.

To determine the positions of nucleosomes reconstituted on YNN1-5, we purified nucleosomes containing these NPEs from the remaining free DNA with sucrose gradient centrifugation. The purified nucleosomes were then subjected to MNase digestion followed by high throughput sequencing (MNase-seq) (**Figure 2B**). To verify the accuracy of nucleosome mapping using this method, we first performed MNase-seq using the 601 nucleosomes. As previously reported (17, 40), our MNase-seq result reveals a single nucleosome position within the 601 DNA when mapping both nucleosome occupancy and dyad distribution (**Figure 2D** and **2E**). The nucleosome coverage of the 601 DNA shows a clear 145bp footprint with a dominant dyad peak that is precisely located at the center of the 145bp 601 sequence.

We then used MNase-seq to determine the position of purified nucleosomes containing YNN1-5 NPEs. MNase-seq measurements of nucleosomes reconstituted with YNN1-5 show distinct patterns in nucleosome occupancy with variable numbers of dyad peaks (**Figure 2D** and **2E**). Nucleosomes reconstituted with YNN2 adopt a single dominant dyad position similar to those on 601. Nucleosomes reconstituted with YNN1 and YNN3 are primarily positioned near the center of the DNA molecules with minor off-center positions. In contrast, nucleosomes reconstituted with YNN4 and YNN5 have a broad number of positions as indicated by both nucleosome occupancy and dyad distribution. These results reveal that nucleosomes reconstituted with YNN1-3, but not YNN4-5, have a well-defined dominate position with different fractions of off-center positions, making them suitable for *in vitro* nucleosome studies.

We further investigated the dyad position of the major YNN1 species by purifying it over an acrylamide gel and performing site-specific crosslinking. As shown in earlier work, site-specific cross-linking histones to DNA, which can be resolved into nicks, can give high-resolution information of nucleosome positions and distributions (36). Cross-links around the nucleosome dyad can be achieved with H3(M120C), where each copy of H3(M120C) nicks DNA both 3’ and 5’ of the dyad. When cross-linking with the purified YNN1 nucleosome was performed, we observed two nicks on each strand, indicating a uniquely positioned histone core (**Figure 3**). Combining information from both strands, the double nicking sites identify the dyad at an AT base pair, where symmetry-related nicks are equidistant in either 5’ or 3’ directions on both strands (**Figure 3C**). This dyad position is 4 bp from that determined by MNase-seq. To further substantiate the dyad position indicated by cross-linking, we also performed site-specific cross-linking with single-cysteine variants of the Chd1 remodeler, which binds symmetrically about the dyad at the two superhelix location 2 (SHL2) sites (31). Cross-links from one variant, Chd1(N459C), are equidistant from and therefore agree with the same dyad position identified by H3(M120C), each giving DNA fragments that migrate between sequencing ladder nucleotides 15-16 on the 5’ side of the dyad of each strand. Cross-links from the other variant, Chd1(V721C), roughly agree with the same dyad, yielding products that migrate between sequencing ladder nucleotides 18-19 on the 5’ side of the dyad on the FAM strand, and between nucleotides 19-20 on the 5’ side of the dyad for the Cy5 strand. These results indicate that MNase-seq on YNN1 determined the dyad position to within four base pairs. This small difference is likely due to the known DNA sequence bias of MNase-seq. Furthermore, these results highlight the importance of carefully characterizing nucleosome positions within physiologically relevant sequences.

**Figure 3.**
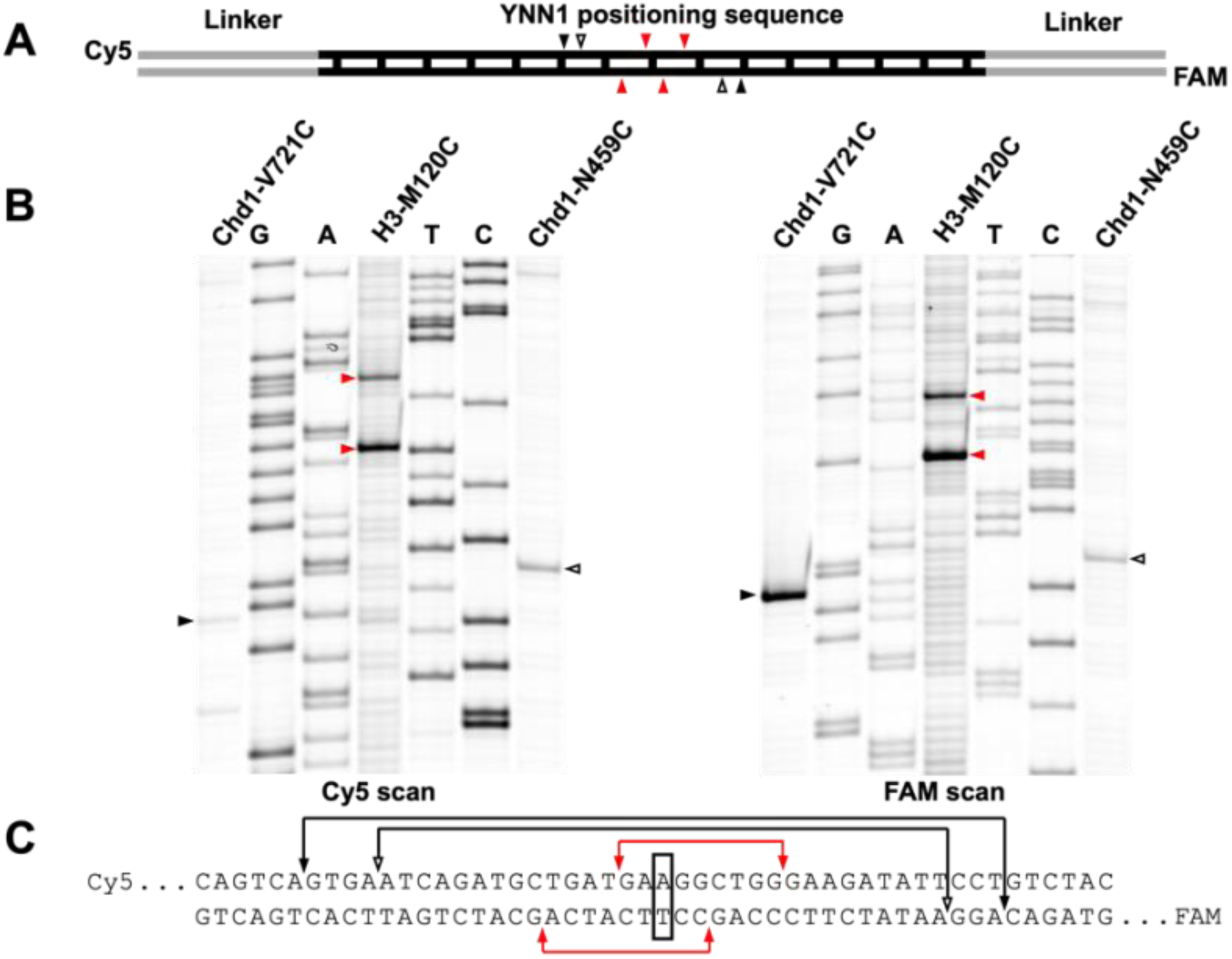
High-resolution mapping of the YNN1 dyad. **(A)** A schematic showing position of crosslinks with respect to the nucleosomal DNA. Crosslinks are shown as closed black, open black, and red arrowheads originating from Chd1-V721C, Chd1-N459C, and H3-M120C, respectively). The vertical black bars represent the SHL divisions with the thickest one representing the dyad. **(B)** Single cysteine mutants of Chd1-N459C (ATPase lobe 1), Chd1-V721C (ATPase lobe 2), and histone H3-M120C were separately labeled with APB and crosslinked to 40-YNN1-40 nucleosomes. Chd1- and H3-DNA crosslinks are visualized by urea denaturing gel and positions determined relative to an enzymatically-generated sequencing ladder. **(C)** Crosslinks are mapped to the YNN1 DNA sequence, and dyad is determined based on the midpoint of the crosslinks on the two strands.

### Nucleosomes with YNN sequences are significantly less stable than with the 601 NPE, yet stability does not correlate with the distributions of nucleosome positions

Based on our observation that the YNN sequences exhibit significantly different nucleosome position distributions, we hypothesized that their free energies (ΔG) of nucleosome formation would vary substantially. Previous work by Widom and co-workers used nucleosome competitive reconstitutions to demonstrate that the 601 NPE has a much lower ΔG than the 5S NPE, suggesting that DNA sequences with lower ΔG form better positioned nucleosomes (17). To test this hypothesis, we used competitive reconstitutions to determine nucleosome formation free energy of YNN1-7 relative to the 5S NPE, which is reported as ΔΔG. Since we determined the location of the nucleosome dyad distributions of YNN1-5 from MNase-seq, we shifted the 247 bp sequences of YNN1-3 into YNN1.c, YNN2.c, and YNN3.c so that the primary dyad positions are located at the center. The YNN4 and YNN5 sequences were kept the same as there is no primary dyad position.

The competitive reconstitution method determines the relative Gibbs free energy (ΔΔG) for nucleosome formation between different DNA sequences (34). In this approach, nucleosome reconstitutions are performed using salt dialysis nucleosome reconstitutions with a DNA molecule containing the NPE of interest, purified histone octamer, and competitor DNA. The competitor DNA is a 147 bp DNA molecule with a sequence from the ampicillin resistance gene of pUC19. The total DNA concentration is set so that only a fraction of DNA is converted into nucleosomes, while the relative concentration of the NPE and competitor DNA is chosen so that the NPE DNA does not fully convert to nucleosomes. During the gradual reduction in electrostatic screening through salt dialysis, the NPE and competitor DNA “compete” for histone octamer binding. The relative concentration of NPE DNA that is wrapped into nucleosomes relative to free NPE DNA is equal to the equilibrium constant, [DNA_nuc_] / [DNA_free_] = K_eq_, which can be determined with EMSAs using native PAGE (34). This equilibrium constant for DNA wrapping into a nucleosome is related to the free energy of nucleosome formation, ΔG, by the Boltzmann distribution through ΔG = -RT ln(K_eq_). By comparing the K_eq_ for each NPE DNA, the ΔΔG between two different DNA sequences is inferred. We followed the convention of using the 5S DNA as the reference DNA (17).

Native PAGE analysis of the competitive reconstitutions with YNN1-7 reveals that they all have slightly lower tendencies to form nucleosomes than the 5S sequence (**Figure 4A**). This is in contrast with the 601 NPE, which reconstitutes into nucleosomes very efficiently. Quantitative analysis for competitive reconstitutions in triplicate reveals that the 601 sequence has a negative ΔΔG of −1.9 kcal/mol (**Figure 4B**, **Table S4**), which implies that the 601 sequence more favorably wraps into nucleosomes than the 5S sequence, as previously reported (17). In contrast, all YNN sequences exhibit positive ΔΔGs ranging between **0.16 and 0.48 kcal/mol** (**Figure 4B**). Importantly, the ΔΔG values for the YNN sequences do not correlate with the level of nucleosome positioning (**Table S4**). For example, YNN2, which reconstitutes into well-positioned nucleosomes (**Figure 2E**), has the most positive ΔΔG value, implying that it wraps the weakest into nucleosomes. Overall, these results reveal that the YNN sequences have similar overall stabilities that are comparable to the native 5S sequence and that the DNA-histone binding free energies do not correlate with the level of nucleosome position homogeneity.

**Figure 4.**
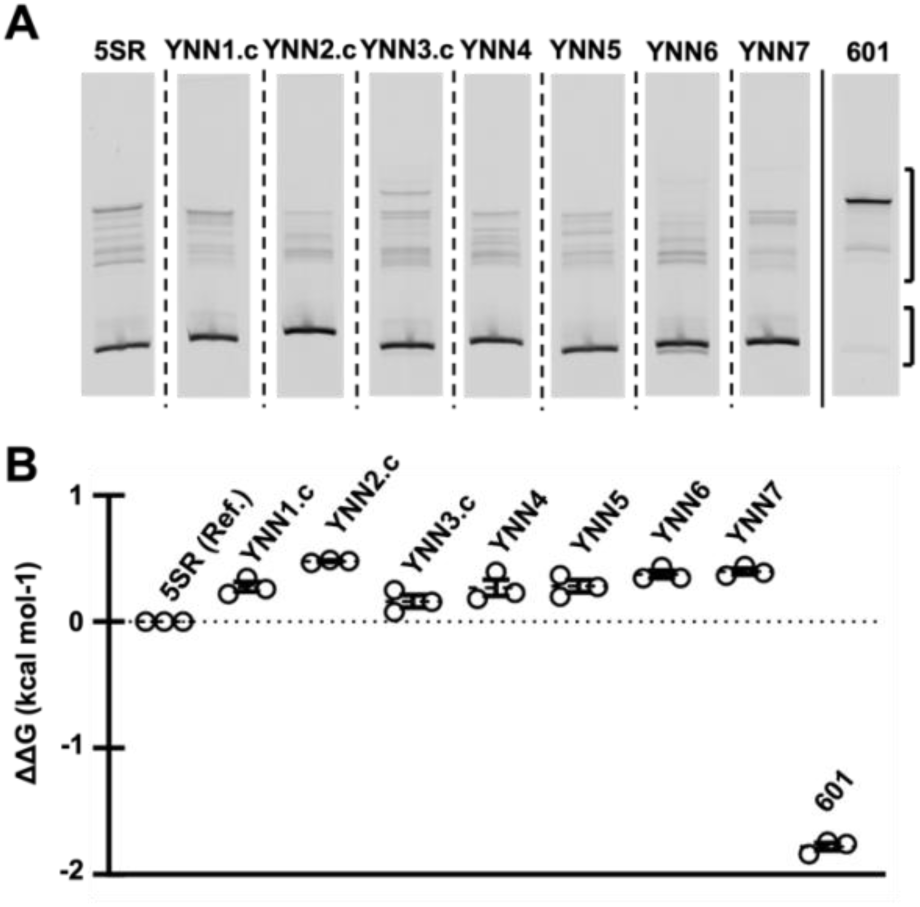
YNNs form nucleosomes that are significantly less stable than the 601 NPE. **(A)** EMSAs that indicate the conversion of the NPE DNA into nucleosomes following competitive reconstitution. Cy3-labeled 247bp DNA was used to reconstitute with non-labeled histone octamers into nucleosomes. The imaging was obtained by detecting the fluorescence of Cy3-labeled DNA. The images of each lane are from the same gel and were rearranged to be in order of the naming convention. **(B)** ΔΔG was determined from the EMSAs of each competitive nucleosome reconstitution. The error bars represent the standard error from three replicates. Since the ΔΔG of 601 NPE and YNN NPEs are standardized from the free energy of 5SR NPE, the ΔΔG of 5SR NPE is 0.

### Transcription factors shift nucleosomes assembled with a native yeast nucleosome positioning element but not with 601-containing nucleosomes

Our observation that YNNs are less stable implies that they may be more mobile than 601 nucleosomes. To test this idea, we investigated if the binding of TFs can directly reposition nucleosomes. Previous studies have revealed that the budding yeast pioneer TFs, Reb1 and Cbf1, are more efficient in displacing nucleosomes than the non-pioneer TF Pho4 *in vivo* (6, 30, 41, 42). *In vitro* studies using 601 nucleosomes show that Reb1 and Cbf1 use a “dissociation rate compensation” mechanism to achieve comparable affinities towards target sites within nucleosomes vs on naked DNA (24). In contrast, Pho4 shows a much lower affinity towards nucleosomes relative to its DNA binding affinity but can still bind to nucleosomes in a partially unwrapped state at high enough concentrations. Importantly, FRET data indicates that TF binding within partially unwrapped nucleosomes does not reposition 601 nucleosomes (24). One difference between the *in vivo* and *in vitro* results could be that the 601 sequence suppresses TF-induced nucleosome repositioning.

To investigate the impact of these TFs on nucleosome repositioning, we prepared the YNN1 and the 601 NPEs with TF motifs that are positioned within the DNA entry-exit region of the nucleosome. We then carried out MNase-seq studies of these nucleosomes with and without a TF. The MNase footprint may change depending on whether the TF is bound and/or if the nucleosome is repositioned. We focused on YNN1 NPE because it has one major position and one minor position separated by 29 bps (**Figure 2**), while using the 601 as a low mobility control. Both sequences contain 50 bps of flanking DNA on each side with a total length of 247 bps (**Figure 5A**), providing additional DNA required for nucleosome repositioning.

**Figure 5.**
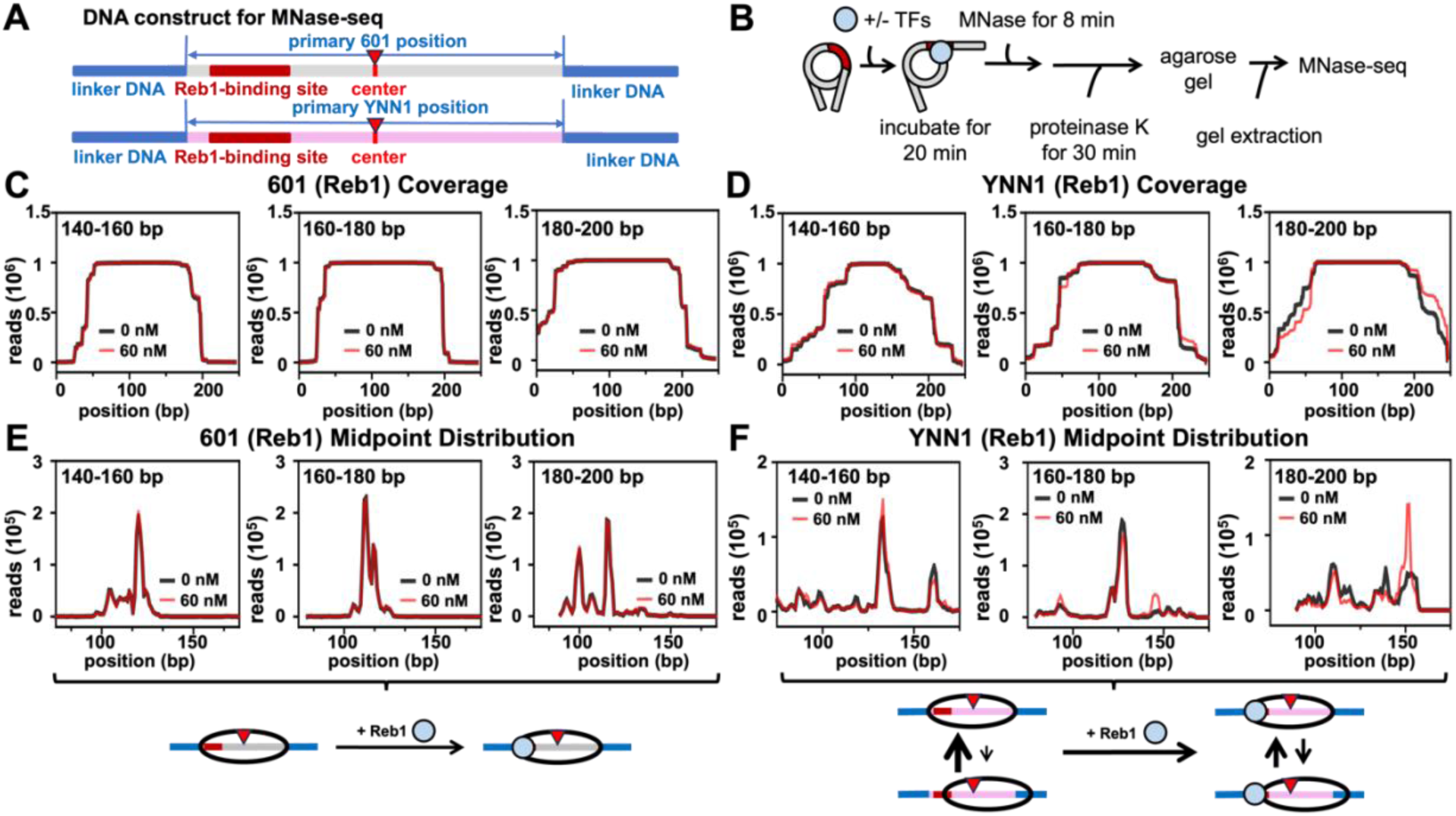
Reb1 repositions YNN1 containing nucleosomes but not nucleosomes with the 601 NPE. **(A)** 601 and YNN1 DNA constructs were used for quantifying the influence of Reb1 binding on nucleosome positions with MNase-seq. 147bp core nucleosome region was labeled as the primary position. The Reb1-binding site (dark red box) was inserted so it starts 8bp into each NPE (**Table S3**). The center of the DNA segment is pointed with a red opposite triangle. **(B)** Workflow for MNase-seq measurements with and without Reb1. The reaction contained 10 nM Cy5-labeled nucleosomes with 0 or 60 nM Reb1. The reaction was incubated at room temperature for 20 minutes and then digested with MNase for 8 minutes at 37 ⁰C. **(C** and **D)** Nucleosome coverage of 601 and YNN1 nucleosomes with (red line) and without (black line) Reb1, respectively. **(E** and **F)** Midpoint distribution of 601 and YNN1 nucleosomes with (red line) and without (black line) Reb1, respectively. The schematics below panels (E) and (F) show the interpretation of the MNase studies.

MNase-seq measurements were carried out with 60 nM of each TF (Reb1, Cbf1, and Pho4) because at this concentration the TF binds a significant fraction of nucleosomes with minimal nucleosome aggregation (**Figure S4**). 60 nM is also roughly equal to the concentration of nucleosomes used in the assay, so the binding here is approximately stoichiometric. To investigate the position of nucleosomes and the footprint of nucleosome plus TF that suppressed MNase cleavage, we categorized the MNase-seq data into three groups based on DNA fragment lengths following MNase digestion: 140-160 bp, 160-180 bp, and 180-200 bp (**Table S5**). Nucleosomes without TF binding predominantly exhibited a 140-160 bp footprint **(46% ± 3%**), consistent with fully wrapped nucleosomes. However, subpopulations with footprints of 160-180 bp (**16% ± 6%**) and 180-200 bp (**1.9% ± 0.7%**) were also observed, likely due to incomplete digestion. TF binding to nucleosomes may shift the distribution of these fragment sizes.

We first investigated the impact of Reb1 on nucleosome positions due to its high affinity towards nucleosomal sites (6, 24) and its frequent *in vivo* localization within the DNA entry-exit region of nucleosomes (43). After incubation with Reb1, we performed MNase-seq to quantify the nucleosome positioning (**Figure 5B, S5**). In the presence of Reb1, nucleosomes that remain in their original position but are partially unwrapped with Reb1 bound will have a 140-160 bp footprint, similar to fully wrapped nucleosomes without Reb1. This similarity is due to Reb1 protection of the unwrapped DNA from MNase digestion. However, nucleosomes that move away from an occupied Reb1 binding site will have larger footprints, with sizes depending on the repositioning distance.

On the 601 NPE, Reb1 binding did not induce a detectable change in the distribution of nucleosome occupancy or dyad position (**Figure 5C and 5E**). These results are consistent with our previous conclusions showing that Reb1 binding does not lead to repositioning of 601 nucleosomes (23, 24, 44). In contrast, Reb1 binding has a significant impact on the nucleosome positioning over the YNN1 NPE. The 140-160 bp length distribution indicates that Reb1 binding modestly increased the fraction of nucleosomes at the primary position, with a corresponding decrease at the minor position **(Figure 5D and 5F**). In contrast, the 160-180 bp and the 180-200 bp length distributions show that Reb1 binding also “pushed” a fraction of nucleosomes from the primary position, increasing the fraction of nucleosomes at the minor position ∼20 bp downstream (**Figure 5D and 5F**). Furthermore, the fraction of fragments in the 180-200 bp group increased 8-fold from 0.4% ± 0.1% to 2.55% ± 0.03% (**Table S5**). These results indicate that nucleosomes containing a native NPE can be repositioned by the pioneer TF Reb1, while nucleosomes with a highly stable NPE like 601 remain resistant to repositioning.

To determine the impact of TFs with different pioneering activities on nucleosome repositioning, we carried out similar analyses using two other yeast TFs, Cbf1 and Pho4 **(Figure S6**). Both TFs contain a basic helix-loop-helix (bHLH) DNA-binding domain, but they differ in their ability to bind nucleosomal DNA. Cbf1, a pioneer TF, binds nucleosomal sites efficiently through dissociation rate compensation, while Pho4 does not exhibit this ability (30). The nucleosome coverage and dyad distributions of the 140-160 bp group showed almost no impact of Cbf1 on nucleosome position, while Pho4 modestly increased the fraction of nucleosomes at the YNN1 major nucleosome positions. In contrast, the 160-180bp and 180-200bp length distribution indicated that both Cbf1 and Pho4 binding within YNN1 nucleosomes induces a shift in nucleosome positions from the central major position to the minor position of YNN1. Moreover, the fraction of the 180bp-200bp length distribution group increased 1.5-fold (from 1.4% ± 0.1% to 1.9% ± 0.4%) after Cbf1 binding and 2.3-fold (from 1.9% ± 0.2% to 4.4% ± 0.4%) after Pho4 binding. These results indicate that nucleosomes wrapped with genomic sequences can be repositioned by either Cbf1 or Pho4, despite their different efficiency of binding within nucleosomes.

### Reb1 targets native or 601 nucleosomes with similar efficiencies

The above results indicate that Reb1 may unwrap and reposition YNN1 nucleosomes. To further corroborate this finding, we utilized FRET efficiency measurements (18, 19)) to probe the structural changes of nucleosomes upon Reb1 binding. We prepared Cy3-Cy5-labeled nucleosomes with DNA that contained (i) the NPE with the Reb1 binding site at the same position used in the MNase-seq measurements and (ii) an additional 60 bp of DNA that extends out from the NPE on the side distal from the Reb1 binding site (**Figure 6A-B**). The Cy5-fluorophore is attached to H2A(K119C), while the Cy3-fluorophore is attached to the DNA at one of two positions: Cy3-proximal and Cy3-distal (**Figure 6A-B**). In the Cy3-proximal construct, Cy3 is positioned to undergo high FRET efficiency with Cy5 in fully wrapped nucleosomes at the NPE position, and so the FRET efficiency drops when nucleosomes are partially unwrapped and/or repositioned from the NPE. In the Cy3-distal construct, Cy3 undergoes intermediate FRET efficiency with Cy5 in fully wrapped nucleosomes at the NPE position, while the FRET efficiency should increase when nucleosomes reposition away from the Reb1 binding site. Importantly, Reb1 binds all 4 of these nucleosome constructs with similar nanomolar apparently dissociation constants based on EMSA measurements (**Figure 6C-D**).

**Figure 6.**
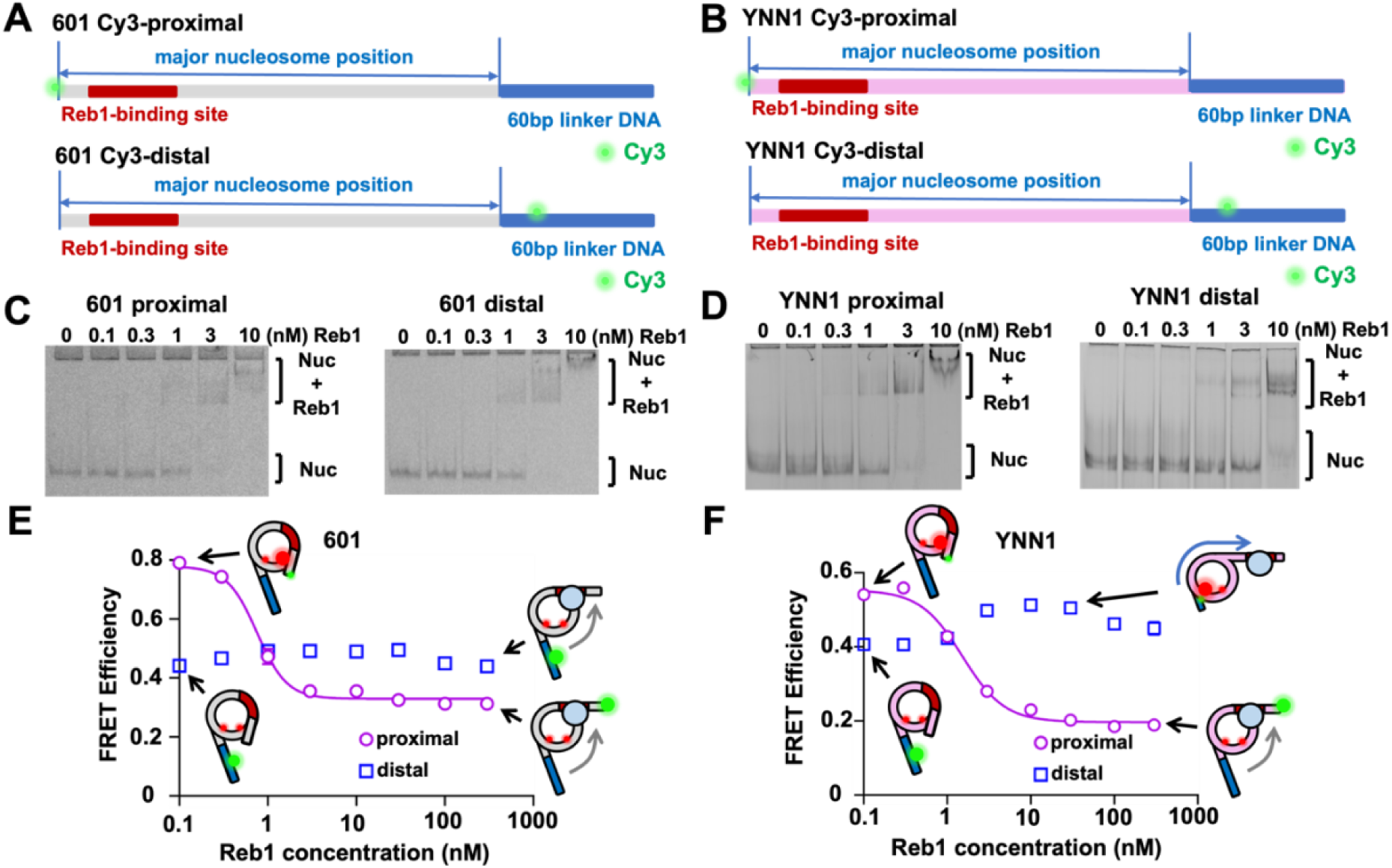
Reb1 targets with similar efficiencies partially unwrapped nucleosomes containing either the YNN1 or the 601 NPE. **(A** and **B)** 601 and YNN1 constructs used in ensemble FRET experiments, respectively. The Cy3-proximal nucleosome constructs position the Cy3 fluorophore near the TF binding site and will undergo high FRET efficiency without a TF (**Table S3**). Both DNA unwrapping and nucleosome repositioning that are trapped by TF binding will reduce the FRET efficiency. The Cy3-distal nucleosome construct positions the Cy3 fluorophore opposite to the TF binding site (**Table S3**), where it undergoes intermediate FRET efficiency with Cy5-labeled histone octamer. TF binding that only traps nucleosomes in partially unwrapped states will not impact the FRET efficiency, while changes in nucleosome position will increase the FRET efficiency. **(C)** EMSAs of Reb1 binding to 601 Cy3-proximal and 601 Cy3- distal nucleosomes. **(D)** EMSAs of Reb1 binding to YNN1 Cy3-Proximal and YNN1 Cy3-distal nucleosomes. **(E)** Ensemble FRET measurements of Reb1 titrations separately with 0.5 nM of 601 Cy3-proximal or 601 Cy3-distal nucleosomes. The reduction in FRET from the 601 Cy3- proximal nucleosomes was fit to a binding isotherm with a S_1/2_ of 0.75 ± 0.04 nM. **(F)** Ensemble FRET measurements of Reb1 titrations separately with 0.5 nM of YNN1 Cy3-proximal of YNN1 Cy3-distal nucleosomes. The reduction in FRET from the YNN1 Cy3-proximal nucleosomes was fit to a binding isotherm with a S_1/2_ of 1.5 ± 0.1 nM.

We first carried out Reb1 titrations with the Cy3-proximal nucleosomes to determine the impact of the NPE on Reb1 binding within partially unwrapped nucleosomes. As previously reported (24), we find that Reb1 binding to 601 Cy3-proximal nucleosomes significantly reduces the FRET efficiency with a half saturation concentration: S_1/2_ = 0.75 ± 0.07 nM (**Figure 6E**, **Table S6**). We then measured Reb1 titrations with YNN1 Cy3-proximal nucleosomes and observed a similar reduction in FRET efficiency with an S_1/2_ = 1.5 ± 0.1 nM (**Figure 6F**, **Table S6**). While this is only 2-fold larger than 601 Cy3-proximal nucleosomes, the difference is statistically significant (P-value = 0.0036). This indicates that the unwrapping probability of YNN1 nucleosomes is about 2-fold less than 601 nucleosomes, implying that the NPE has a modest but quantitative impact on TF occupancy within partially unwrapped nucleosomes, which is consistent with previous studies that compared 5S and 601 NPEs (45).

We then carried out Reb1 titrations with the Cy3-distal nucleosomes to determine if the FRET efficiency can detect Reb1-induced nucleosome repositioning. The Reb1 binding to 601 Cy3-distal nucleosomes increased the FRET efficiency from 0.440 ± 0.004 to 0.488 ± 0.005 at 10 nM Reb1, the concentration where nucleosomes are fully bound (**Figure 6E**, **Table S6**). While the difference in FRET efficiency is only 0.048 ± 0.006, this difference is statistically significant (P-value = 0.0017). Reb1 binding to YNN1 Cy3-distal nucleosomes increased the FRET efficiency from 0.407 ± 0.005 at 0.1 nM Reb1 to 0.513 ± 0.006 at 10 nM Reb1 (**Figure 6E**, **Table S6**). This increase in FRET efficiency of 0.106 ± 0.008 is statistically significant (P-value = 0.0002) and over 2-fold larger than with 601 nucleosomes. These results indicate that Reb1 binding repositioned YNN1 nucleosome more efficiently than 601 nucleosomes, which confirms our MNase-seq measurements. Furthermore, the combination of the Cy3-proximal and Cy3-distal measurements is consistent with previous studies that showed that the central ∼80 bps of the 601 NPE are fully responsible for the strong DNA-histone affinity (46) and that the outside ∼35 bps of the 601 sequence partially unwrap with similar probabilities as the 5S NPE (45).

### The rate of nucleosome sliding by Chd1 is influenced by native NPE sequences

For ATP-dependent remodelers, repositioning is often measured by tracking nucleosome migration in native gels over time using quantitative EMSA analysis (47). For the Chd1 remodeler, DNA sequence can affect the rate of nucleosome sliding where end-positioned nucleosomes with flanking DNA on either side of the 601 sequence differ by ∼6-fold in sliding rates (26). Compared to the 601 NPE, the YNN NPEs provide the opportunity to measure remodeling rates on nucleosomes with lower stabilities and a variety of DNA sequences. We focused on the highly conserved chromatin remodeling complex, Chd1 (48–50), a member of the CHD family that establishes and maintains evenly spaced nucleosomes within chromatin (51–53).

To investigate Chd1 remodeling, we relied on its ability to preferentially shift end-positioned mononucleosomes away from short linker DNA and toward longer linker DNA, a characteristic believed to underlie the generation of evenly spaced arrays (54, 55). Briefly, we reconstituted pairs of end-positioned nucleosomes with 207 bps DNA fragments containing 601 or YNN 1-4 NPEs, with flanking DNA either on only the right or left side of the positioning sequence (**Figure 7A**). These nucleosomes (10 nM) were incubated with Chd1 (10 nM) and ATP (1 mM), and their repositioning over time was analyzed using EMSA assays. As previously reported, Chd1 repositions nucleosomes away from the end-positioned NPEs towards more central positions within the DNA (**Figure 7B**). For 601 nucleosomes, the rates of Chd1 remodeling in the two directions differ by a factor of ∼4 (*k_fast_* = 0.31 ± 0.04 min^-1^, *k_slow_* 0.08 ± 0.01 min^-1^; **Figure 7B-D, S7-S9, Table S7**), consistent with a previous study (26). In addition, 53 ± 2 % and 60 ± 2 % of end-positioned nucleosomes are shifted toward a center position for the slow and fast directions, respectively (**Figure 7E, Table S8**).

**Figure 7.**
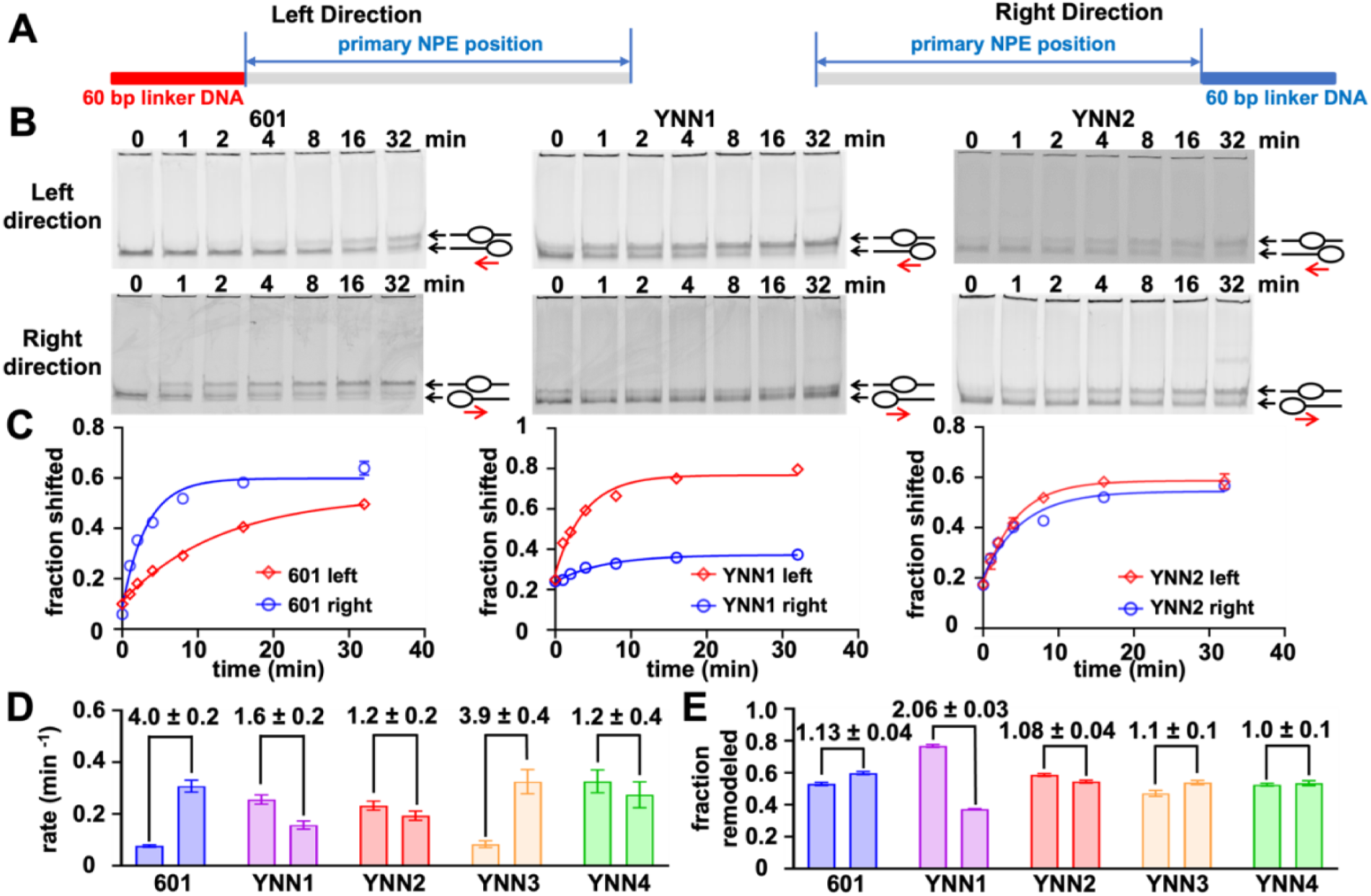
NPE DNA sequence influences the nucleosome remodeling efficiency of Chd1. **(A)** NPE constructs used in nucleosome remodeling experiments. Left Direction has a 60 bp linker on the left side of the NPE, while Right Direction has a 60 bp linker on the right side of the NPE. **(B)** EMSA measurements of Chd1 nucleosome remodeling time courses. 10 nM nucleosomes were incubated with 10 nM Chd1 for 0, 1, 2, 4, 8, 16, 32 minutes. Nucleosomes reposition from end positions to center positions, which results in a change in electrophoretic mobility. **(C)** Quantification of the rate of remodeling and the fraction of nucleosomes that were remodeled by Chd1. Each EMSA measurement was repeated three times and quantified with Image J. Uncertainty bars (which are sometimes smaller than the data point size). The two directions of each NPE were organized based on the left and right directions. **(D)** Bar plot summary of the remodeling rate by Cbf1 of nucleosomes with different NPEs. NPEs were organized based on left and right directions. **(E)** Bar plot summary of the fraction of nucleosomes remodeled by Chd1 of nucleosomes with different NPEs. Different NPEs were organized based on the order of left and right directions.

We next determined the rate and fraction of Chd1 repositioning for nucleosomes reconstituted with YNN1-4. Given their differences in stability, we initially hypothesized that YNN nucleosomes would be remodeled more rapidly. Surprisingly, the rates of remodeling for all four YNN NPEs fall within the range of the fast and slow rates observed for 601 nucleosomes (**Figure 7D, Table S7**). This suggests that nucleosomes with native NPEs are remodeled at similar rates to 601 nucleosomes. The remodeling rates for two of the YNN NPEs, YNN1 and YNN3, are direction-dependent (**Figure 7C, 7D, S7B**). In addition, the fraction of remodeled nucleosomes for all NPEs shows no difference in both remodeling directions except for YNN1, which differs by 2-fold (**Figure 7E, Table S8**).

Given that Chd1 can remodel nucleosomes from an NPE in the two directions at different rates, we considered the possibility that this results in a preferred remodeling direction when the nucleosomes can be remodeled in both directions. To investigate this, we carried out Chd1 remodeling of nucleosomes at the major position of YNN1 with 40 base pairs of linker DNA on each side of the nucleosome, allowing the nucleosomes to be repositioned in either direction. Using site-direct DNA cross-linking described above, we mapped nucleosome positions with H3(M120C) and H2B(S53C) before and after remodeling. We find that at the starting nucleosome position, cross-links were observed for the dyad proximal H3(M120C) but not for H2B(S53C), which should cross-link around SHL5. Upon the addition of Chd1 and ATP, cross-links corresponding to H3(M120C) shift toward the Cy5 (left) side of the YNN1 NPE, and new strong cross-links appear ∼35-45 nt (FAM side, right) and ∼63 nt (Cy5 side, left) from the dyad, which we attribute to H2B(S53C) (**Figure S10**). The inability to initially observe cross-links from H2B(S53C) is likely due to DNA sequence, since the azido-phenacyl bromide cross-linker does not react with thymines (56), and there are AT-rich stretches around SHL5 of the starting YNN1 nucleosome position. Judging by the change in H3(M120C) cross-links, we estimate that Chd1 shifts these nucleosomes ∼10-20 bp. This nucleosome shift corresponds to the fast direction (**Figure 7**). In agreement with previous work showing biased sliding of 601 nucleosomes (20, 56), these results suggest that the asymmetry in sequence-dependent sliding rates bias the steady state positions of remodeled nucleosomes. Overall, these results suggest that, while native NPE sequences can influence the remodeling rate and fraction of nucleosomes remodeled by Chd1, the overall differences are not substantial compared to 601-containing nucleosomes.

## DISCUSSION

In this study, we used biochemical, high-throughput sequencing, and fluorescence assays to characterize a collection of native NPEs from budding yeast, and then utilize them to investigate nucleosome repositioning. Our approach to using both *in vivo* and *in vitro* genome-wide nucleosome positioning data allows us to identify multiple native NPEs. However, the distribution of nucleosome positions within each of the selected sequences varies significantly. Two out of the seven selected sequences do not position nucleosomes. Furthermore, the distribution of nucleosome positioning varied significantly among the five sequences with a dominant nucleosome position. These results highlight the need to carefully characterize any NPE candidate before using it to prepare nucleosomes for biochemical and/or biophysical measurements.

Our studies determined the differences in partial unwrapping, ATP-independent repositioning, and ATP-dependent repositioning between 601 and YNN-containing nucleosomes. We found that partial unwrapping and ATP-dependent repositioning are affected by NPE sequences but are overall similar between 601 and YNN nucleosomes. In contrast, repositioning by TF binding does not occur for 601 nucleosomes, while YNNs can be shifted upon TF binding. These results highlight that high-affinity NPEs such as the 601 are important reagents for the investigation of nucleosome dynamics, but they have limitations. Namely, while 601 appears to work well for nucleosome remodeling by Chd1 and nucleosome unwrapping, studies involving ATP-independent nucleosome repositioning should avoid using 601-containing nucleosomes and instead use native NPEs.

Previous studies have relied on native NPEs for investigating nucleosome dynamics. The most frequently used native NPE is the 5SR sequence (57), which positions nucleosomes similarly to the YNN1 sequence in this study. More recent studies have investigated nucleosome interactions with TFs with native sequences, including PU.1 and Oct4 interacting with nucleosomes containing sequences from the CX3CR1 and LIN28B gene, respectively (25, 58). Interestingly, PU.1 and Oct4 have been reported to reposition nucleosomes containing these native sequences. However, another study showed that Oct4 traps nucleosomes in an unwrapped state instead of shifting the nucleosome position (59, 60). In addition, these results are in contrast to the report that Oct4 targets its site within 601-containing nucleosomes without nucleosome unwrapping or repositioning (61, 62). These previously published findings, combined with our results, highlight how the underlying DNA sequence influences the impact of TF binding on nucleosome partial unwrapping and repositioning.

The 601 NPE was selected based on its preference to form nucleosomes by salt dialysis reconstitution (17). Because the H3-H4 histone tetramer binds DNA before the H2A-H2B heterodimers during these reconstitutions, the selection of 601 was based on DNA interactions with the H3-H4 tetramer. This resulted in the strong DNA-histone interactions being localized to the central 71 base pairs of the 601 sequence (46). This explains why the apparent K_D_ for TF binding within partially unwrapped nucleosomes is similar between YNN1-containing and 601-containing nucleosomes.

The 601 sequence forms a narrow distribution of nucleosome positions due to the high affinity for the H3-H4 tetramer (46). Interestingly, nucleosome position homogeneity for the different YNN NPEs did not correlate with their relative ΔΔG for nucleosome formation. This suggests that the mechanism behind how the YNN NPEs position nucleosomes is distinct from how the 601 NPE positions nucleosomes. One potential explanation is that the homogeneity/sharpness of nucleosome distributions stems from the relative ΔΔG for nucleosome formation between different positions within a specific YNN NPE. This would imply that the DNA sequences outside YNN1 and YNN2 disfavor being wrapped into nucleosomes relative to the central 147 bp YNN1 and YNN2 sequence. This is similar to the known impact of poly-A tracts in gene promoters, which disfavor being wrapped into nucleosomes (63). Future measurements will be required to directly test this hypothesis.

Most work looking at how DNA sequence influences sliding rates has been done with the Widom 601. Surprisingly, the rates of nucleosome sliding by Chd1 that we report here are similar for 601 nucleosomes and native NPE-containing nucleosome, despite the YNN sequences having significantly more positive ΔΔG of nucleosome formation relative to 601 (**Figure 4**). The 601 sequence was shown to have a marked asymmetry in the direction of nucleosome sliding (20, 26, 56), and our work here extends this phenomenon to also include natural sequences. This indicates that remodeling studies (at least with Chd1) with 601-containing nucleosomes provide an accurate picture of how nucleosomes are remodeled with native NPEs. More importantly, this suggests that DNA sequence-dependent remodeling asymmetry could bias nucleosome position distributions *in vivo.* Given the different patterns of nucleosomes generated by different remodeler types (64), it will be important to better understand the sequence dependence of nucleosome remodeling and how it influences not only Chd1 remodeling but also the different families of nucleosome chromatin remodelers.

## DATA AVAILABILITY

Sequencing data has been deposited in the Gene Expression Omnibus database with the accession number GSE287238. The additional experimental data sets are either included in this manuscript, the supplemental information, or are available from the authors upon request.

## ACKNOWLEDGEMENTS

We are grateful to the Bai Lab members, Bowman Lab members, and the Poirier Lab members, especially Michael Neuhoff, for insightful discussions and feedback on this work. We acknowledge Cheryl Keller of the Genomic Research Incubator at Penn State, for technical support. This work was funded by the National Institutes of Health [R35 GM139564 to M.G.P., R35 GM139654 to L.B., and R01 GM084192 to G.D.B.]; National Science Foundation [MCB 2411725 to M.G.P.]; Funding for open access charge: National Institutes of Health R35 GM139564.

## Supplemental Information

### Supplemental Figures

**Figure S1.**
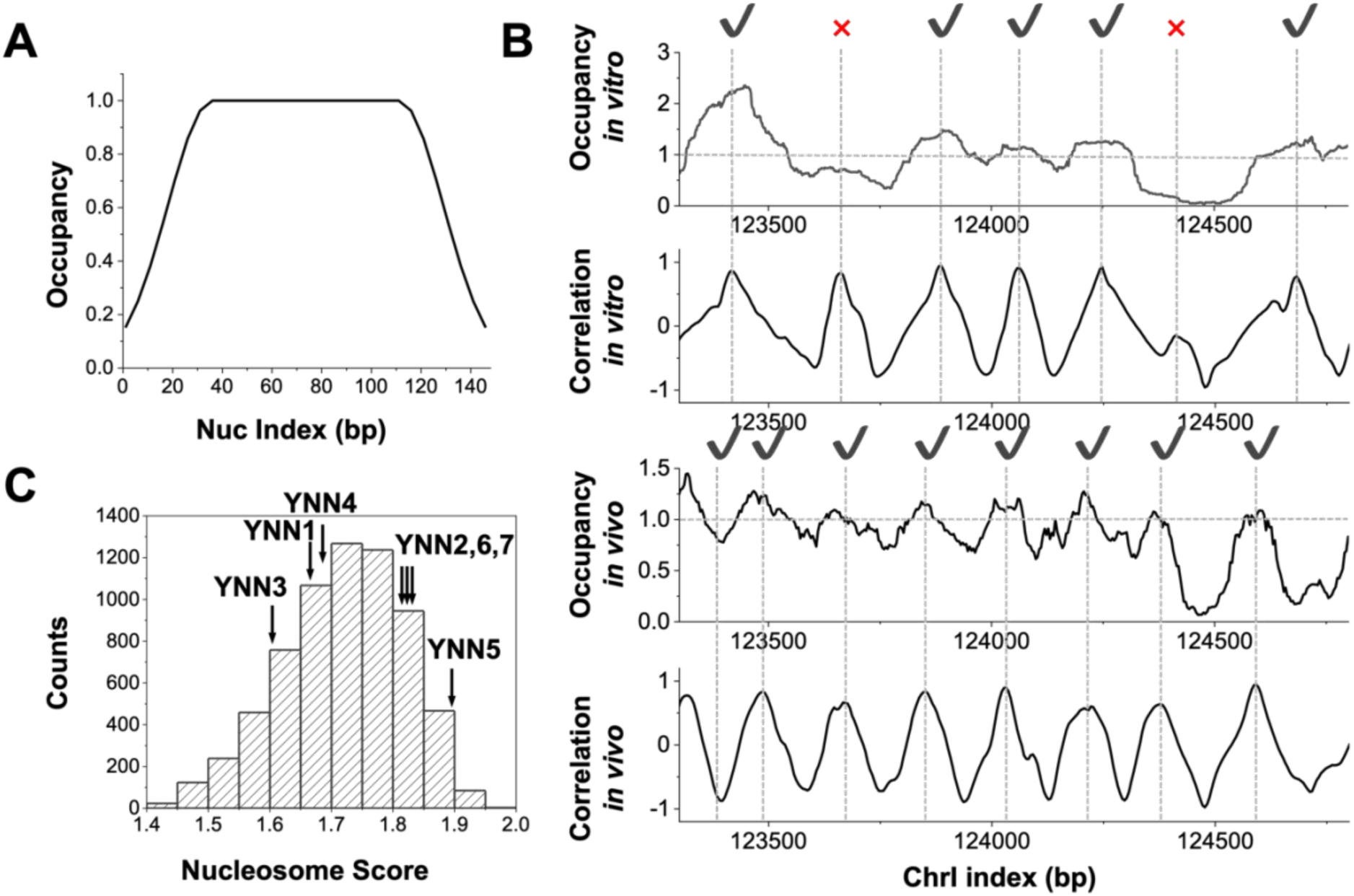
Genome-wide identification of NPE candidates. **A)** Standard nucleosome occupancy profile, representing protection levels against MNase digestion for a well-positioned nucleosome. Out of the 147 bp, the central 80 bp are fully protected, while the entry and exit regions show partial protection, modeled here with a Gaussian function. **B)** Nucleosome occupancy measured *in vitro* and *in vivo*, along with their correlations with the standard nucleosome profile in (A). Positive peaks in the correlation curve with corresponding occupancy values greater than 1 are identified as nucleosome dyads (indicated with a check mark), while occupancy values less than 1 are rejected as nucleosome dyads (indicated with a red x). **C)** Distribution of the scores for aligned nucleosomes identified throughout the genome. The scores are calculated as the sum of their correlation values *in vivo* and *in vitro*. Scores for YNN1-7 are indicated by arrows, illustrating their broad range within this distribution.

**Figure S2.**
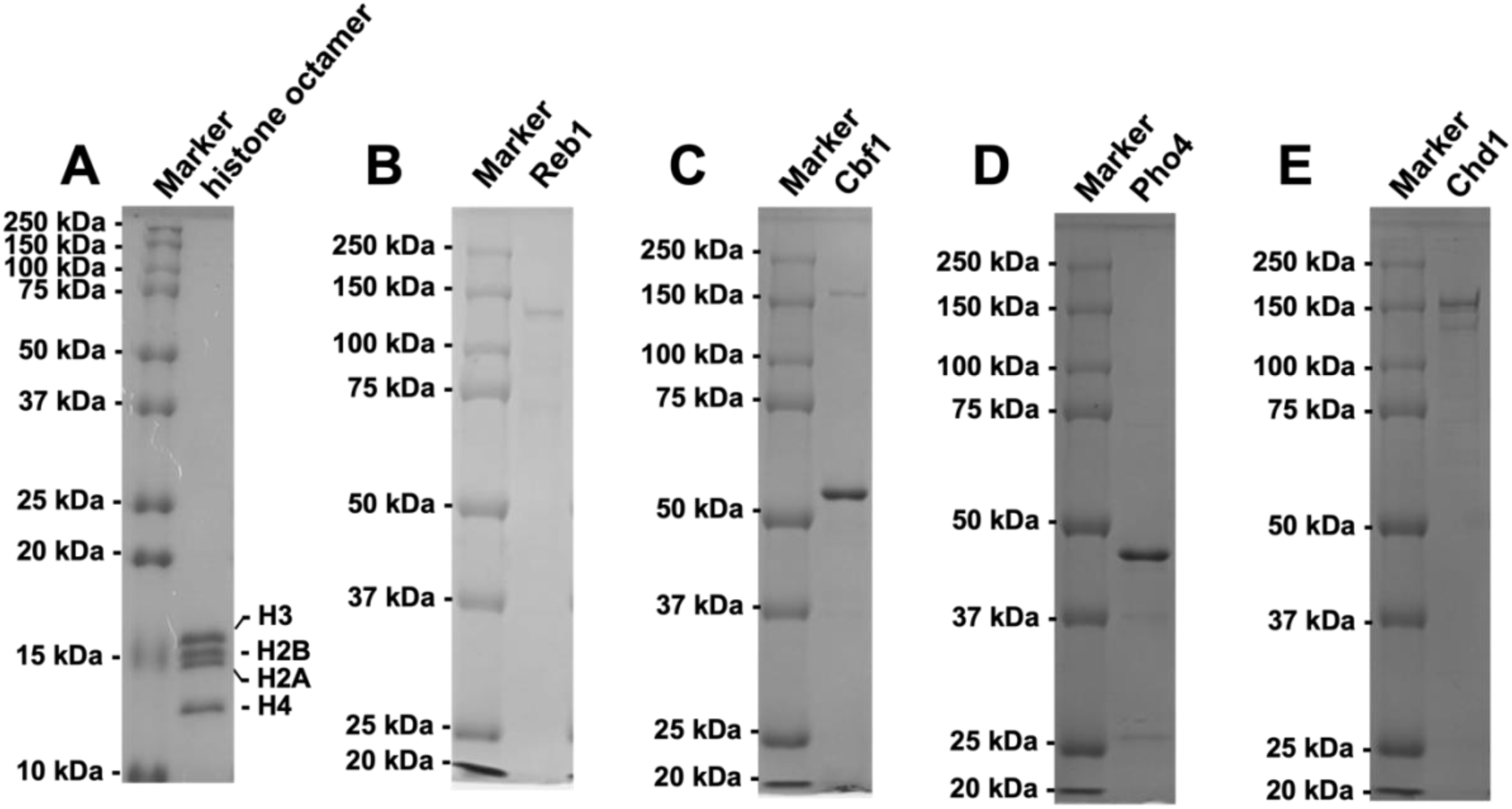
Protein preparation for nucleosome binding and sliding experiments. SDS-PAGE analysis of purified recombinant (**A**) histone octamer, (**B**) Reb1, (**C**) Cbf1, (**D**) Pho4, and (**E**) Chd1. The left lane contains a molecular weight standard, while the right lane contains the protein sample. Each gel is visualized by Coomassie brilliant blue staining.

**Figure S3.**
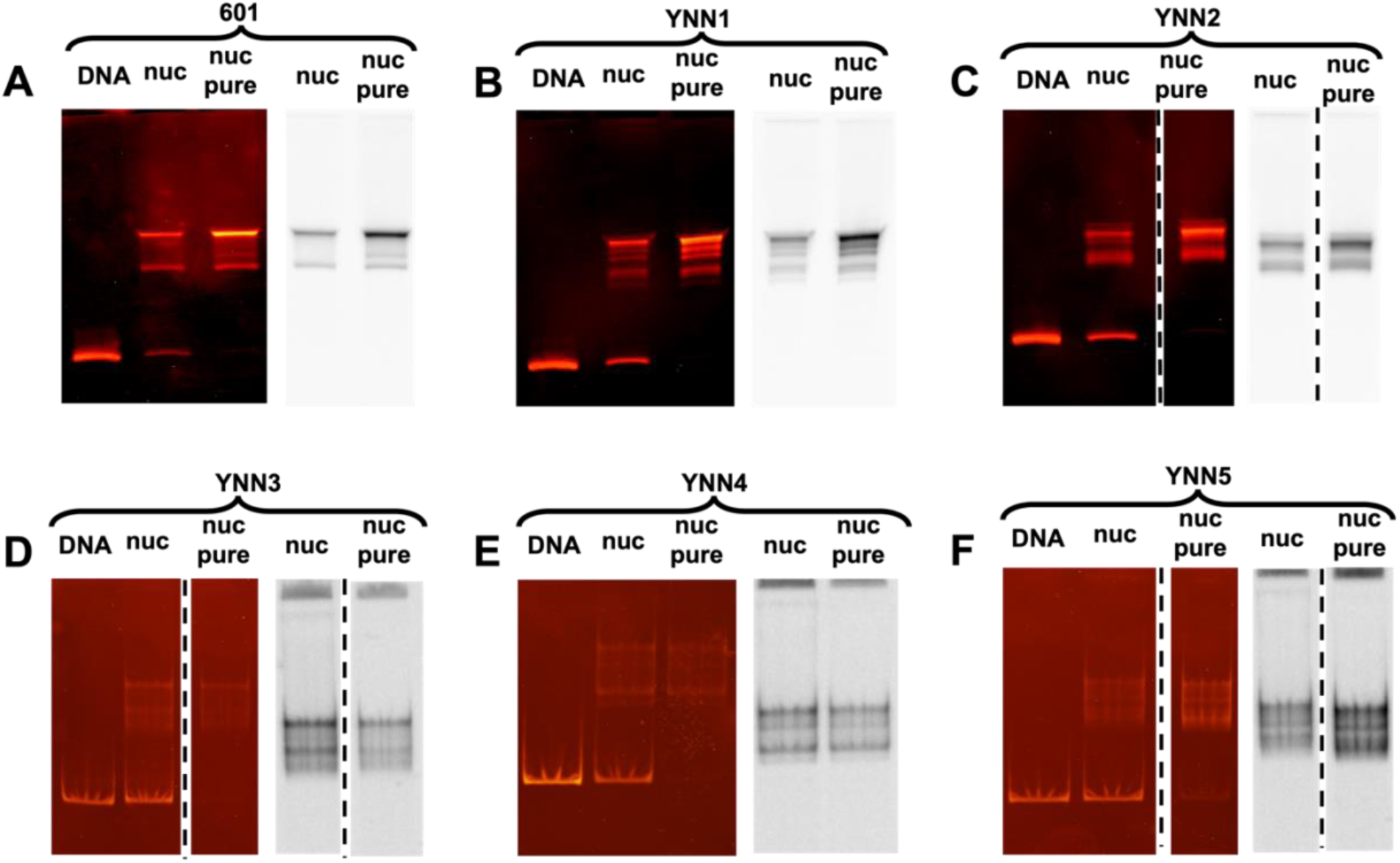
Nucleosome reconstitution of YNNs and 601 nucleosomes. **(A-F)** EMSA of the nucleosomes prepared for MNase-seq studies. Nucleosomes were reconstituted with Cy5- labeled histone octamer. The Cy5 fluorescence scan is on the right, while the ethidium bromide fluorescence image is on the left. DNA lanes contain HPLC-purified naked DNA that was used for nucleosome reconstitution. “nuc” lanes contain the product right after nucleosome reconstitution and before sucrose gradient centrifugation. “nuc pure” lanes contain sucrose gradient purified nucleosomes.

**Figure S4.**
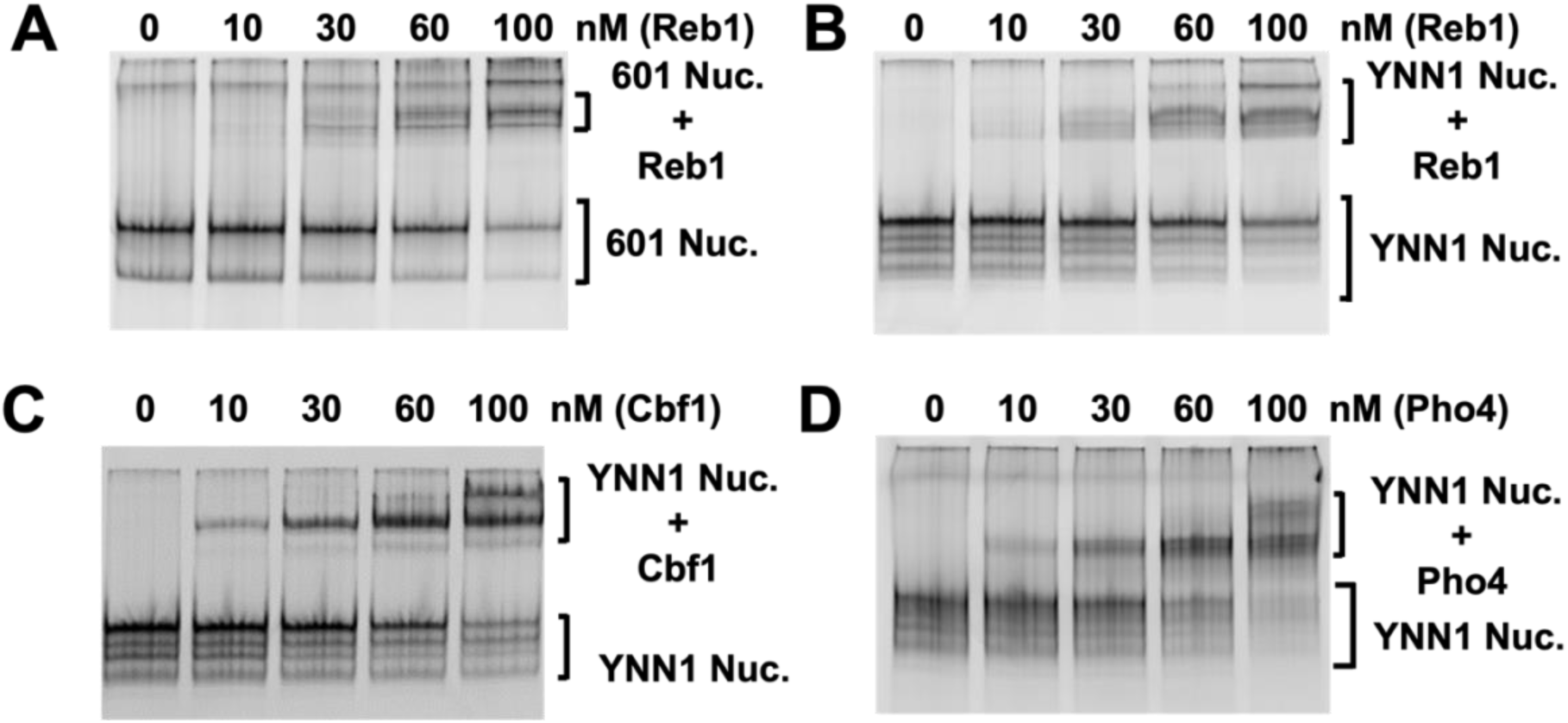
EMSA measurements of TF binding to nucleosomes for MNase-seq studies. **(A)** EMSA measurements of Reb1 binding to 601-containing nucleosomes that are labeled with Cy5 at H2A(K119C). The image is a Cy5 fluorescence scan. The nucleosomes included a Reb1 binding site in the DNA entry-exit region and were at a concentration of 60 nM. **(B)** EMSA measurements of Reb1 binding to YNN1-containing nucleosomes that are labeled with Cy5 at H2A(K119C). The image is a Cy5 fluorescence scan. The nucleosomes included a Reb1 binding site in the DNA entry-exit region and were at a concentration of 60 nM. **(C)** EMSA measurements of Cbf1 binding to YNN1- containing nucleosomes that are labeled with Cy5 at H2A(K119C). The image is a Cy5 fluorescence scan. The nucleosomes included a Cbf1 binding site in the DNA entry-exit region and were at a concentration of 60 nM. **(D)** EMSA measurements of Cbf1 binding to YNN1-containing nucleosomes that are labeled with Cy5 at H2A(K119C). The image is a Cy5 fluorescence scan. The nucleosomes included a Cbf1 binding site in the DNA entry-exit region and were at a concentration of 60 nM.

**Figure S5.**
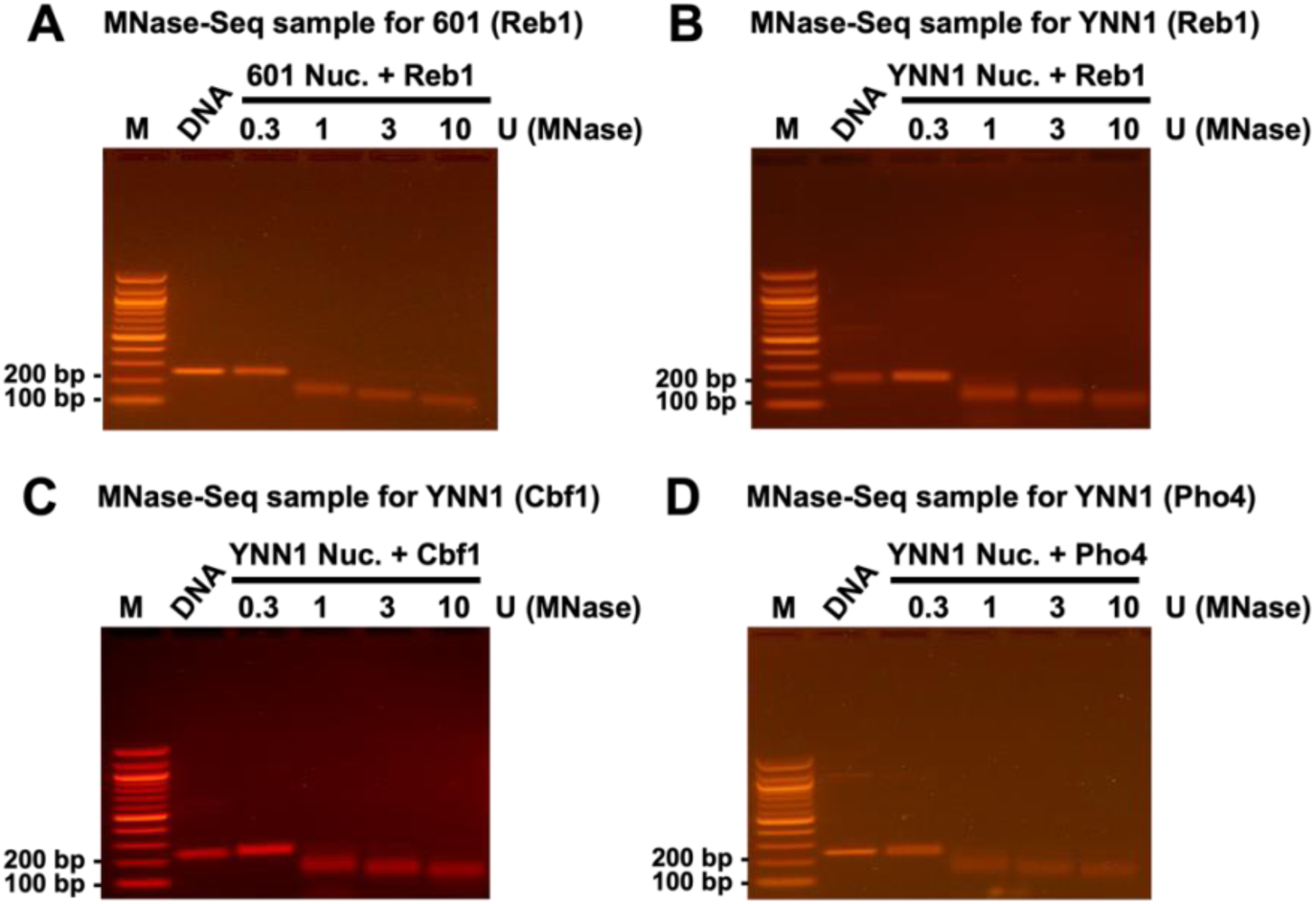
MNase digestions for the MNase-seq measurements. Ethidium bromide fluorescence images of nucleosomal DNA following 8 minutes MNase digestions at 37 ⁰C. 60 nM of a TF was added before the MNase digestion. The nucleosomes studied were **(A)** 601 nucleosomes with Reb1, (**B**) YNN1 nucleosomes with Reb1, (**C**) YNN1 nucleosomes with Cbf1, and (**D**) YNN1 nucleosomes with Pho4. Each lane is labeled with the unit amount of MNase used in the 10 μl reaction volume.

**Figure S6.**
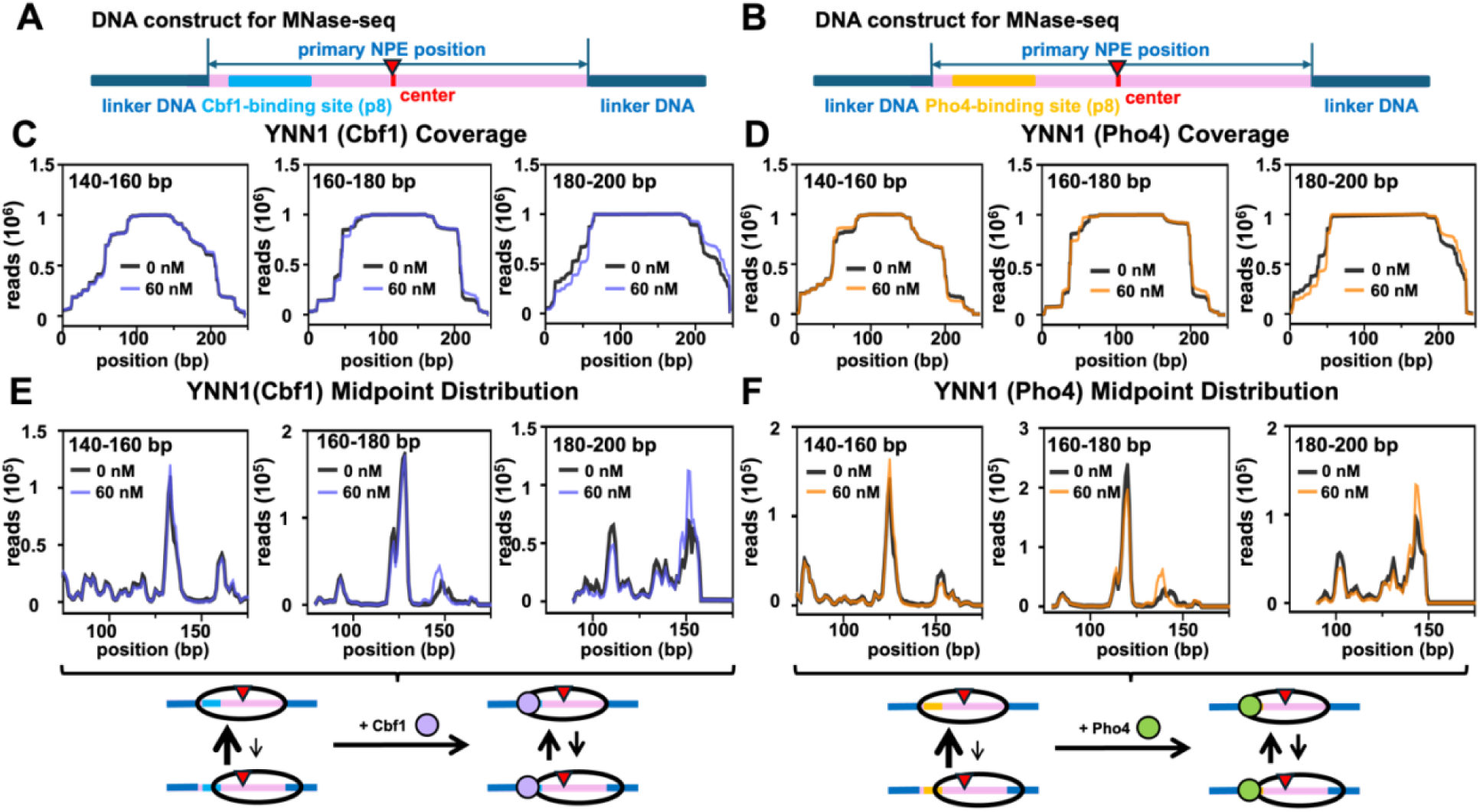
Cbf1 and Pho4 influence YNN1 containing nucleosome positions but not nucleosomes with the 601 NPE. **(A** and **B)** YNN1 DNA constructs used for quantifying the influence of Cbf1 and Pho4 binding on nucleosome positions with MNase-seq. 147bp core nucleosome region was labeled as the primary position. The Cbf1 and Pho4 binding sites (blue and yellow box, respectively) were inserted so they start 8bp into the YNN1 NPE. The center of the DNA segment is pointed with a red opposite triangle. **(C** and **D)** Nucleosome coverage of YNN1 nucleosomes with (red line) and without (black line) either Cbf1 or Pho4, respectively. **(E** and **F)** Midpoint distribution of YNN1 nucleosomes with (red line) and without (black line) either Cbf1 or Pho4, respectively. The schematics below panels (E) and (F) show the interpretation of the MNase studies.

**Figure S7.**
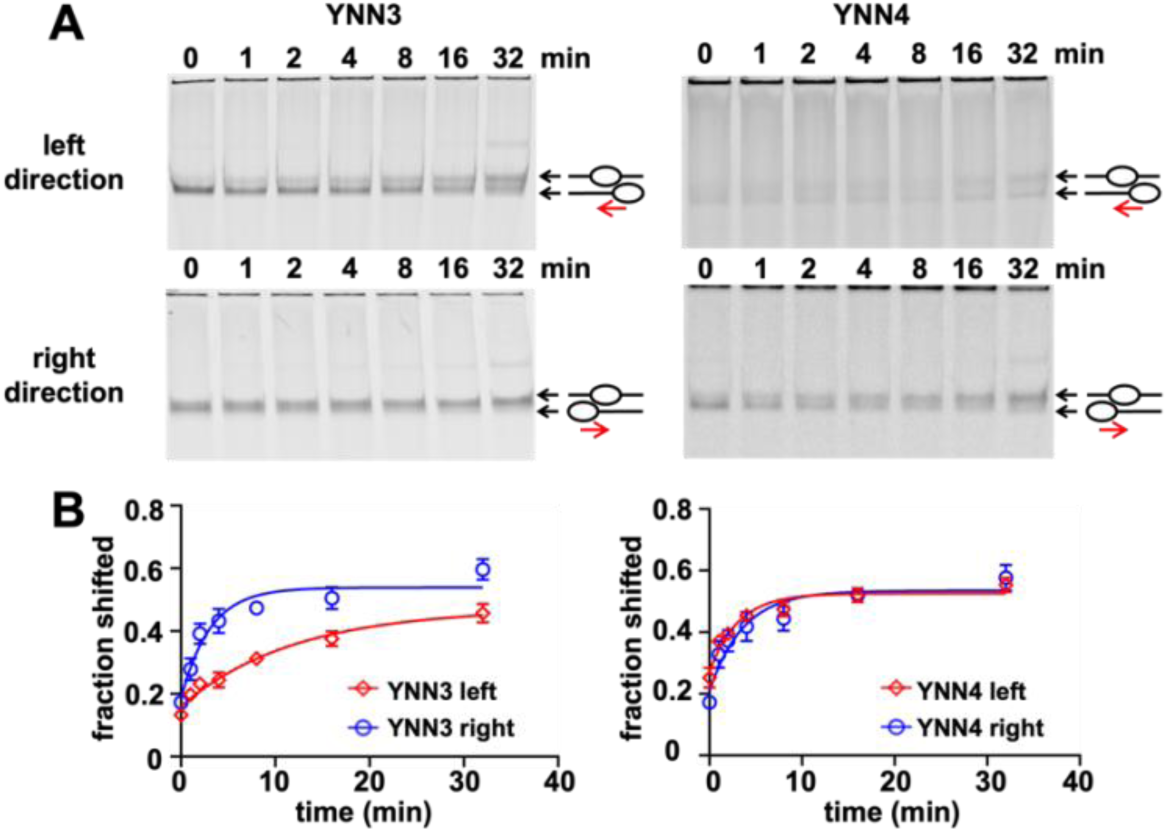
EMSA measurements of Chd1 remodeling of YNN3 and YNN4 nucleosomes. **(A)** EMSA measurements of Chd1 nucleosome remodeling time courses. 10 nM nucleosomes were incubated with 10 nM Chd1 for 0, 1, 2, 4, 8, 16, 32 minutes. Nucleosomes reposition from end positions to center positions, which results in a change in electrophoretic mobility. Each gel is labeled with YNN sequence used in the measurement. **(B)** Quantification of the rate of remodeling and the fraction of nucleosomes remodeled by Chd1. Each EMSA measurement was repeated three times and quantified with Image J. Uncertainty bars (some are sometimes smaller than the size of the data point) represent the standard error of the 3 measurements. The two directions of each NPE were organized based on the left and right directions.

**Figure S8.**
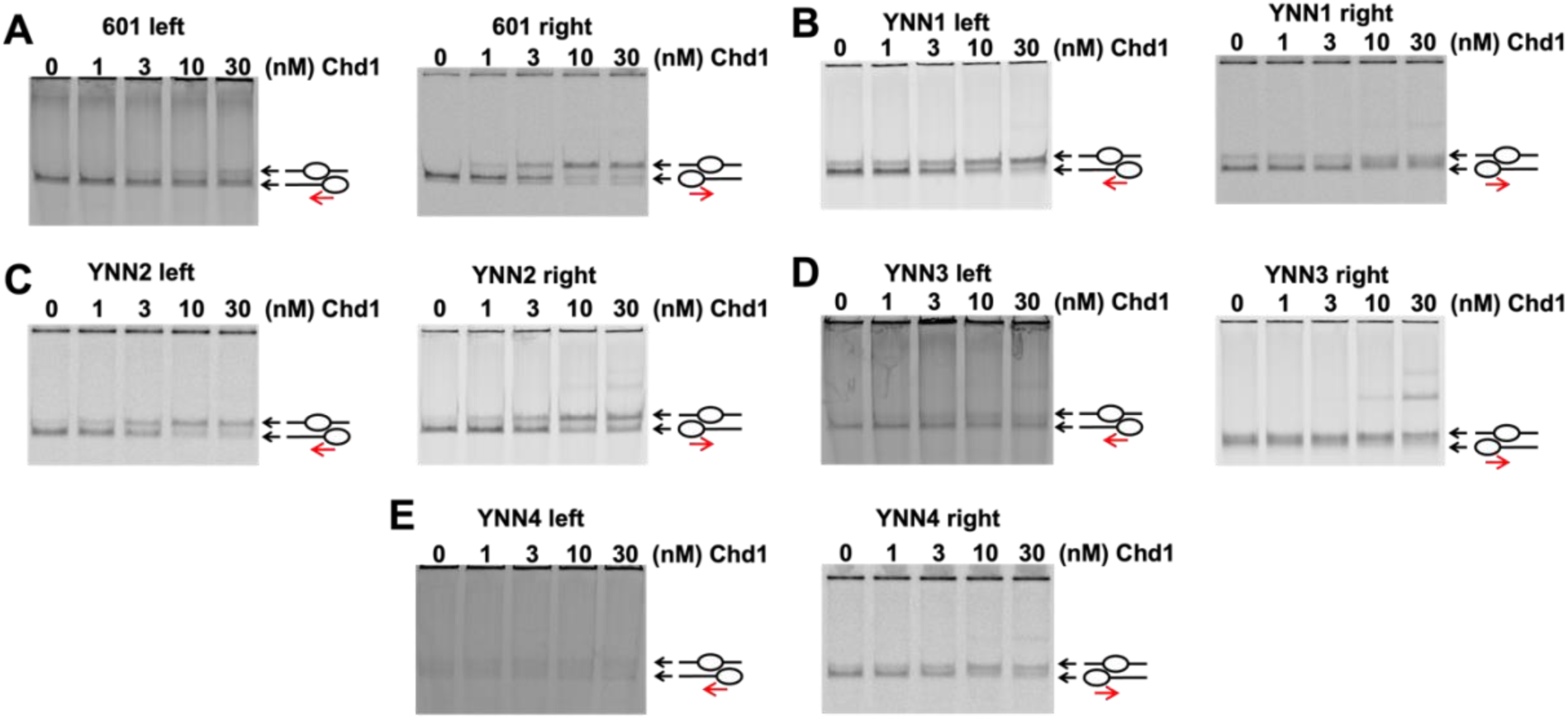
Nucleosome remodeling at different Chd1 concentrations. EMSA measurements of nucleosome remodeling at increasing concentrations of Chd1. Each image is a Cy5 fluorescence scan. The lanes are labeled by the Chd1 concentrations. The measurements were done with 10 nM of nucleosomes that were labeled with Cy5 at H2A(K119C). The EMSA measurements were done with nucleosomes containing (**A**) 601, (**B**) YNN1, (**C**) YNN2, (**D**) YNN3, and (**E**) YNN4. The images were labeled with left or right to indicate if the remodeling direction was in the left or right direction as determined in the remodeling rate measurements (Figure 7).

**Figure S9.**
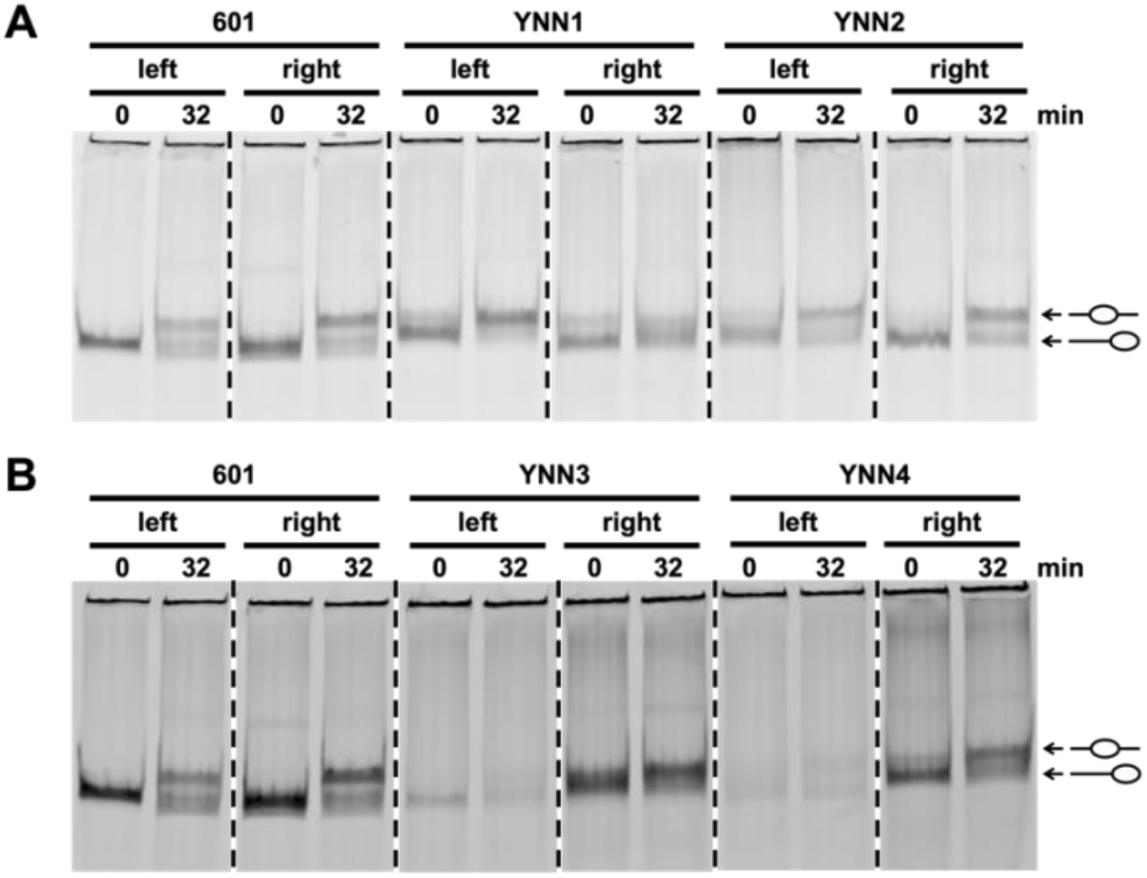
Comparison of the electrophoretic mobility of nucleosomes with different NPEs before and after remodeling by Chd1. EMSA measurements of nucleosome remodeling in the left and right directions. The remodeling reactions were done for 32 minutes with 10 nM of nucleosomes that were labeled with Cy5 at H2A(K119C), and 10 nM Chd1. The lanes are labeled with 0 or 32 to indicate if the lane contains nucleosomes before or after 32 minutes of remodeling by Chd1. (**A**) Cy5 fluorescence scan of the gel that analyzed nucleosomes with the 601, YNN1, and YNN2 sequences. All nucleosomes were loaded in the same gel and re-organized based on left or right directions. (**B**) Cy5 fluorescence scan of the gel that analyzed nucleosomes with the 601, YNN3, and YNN4 sequences. All nucleosomes were loaded in the same gel and re-organized based on left or right directions.

**Figure S10.**
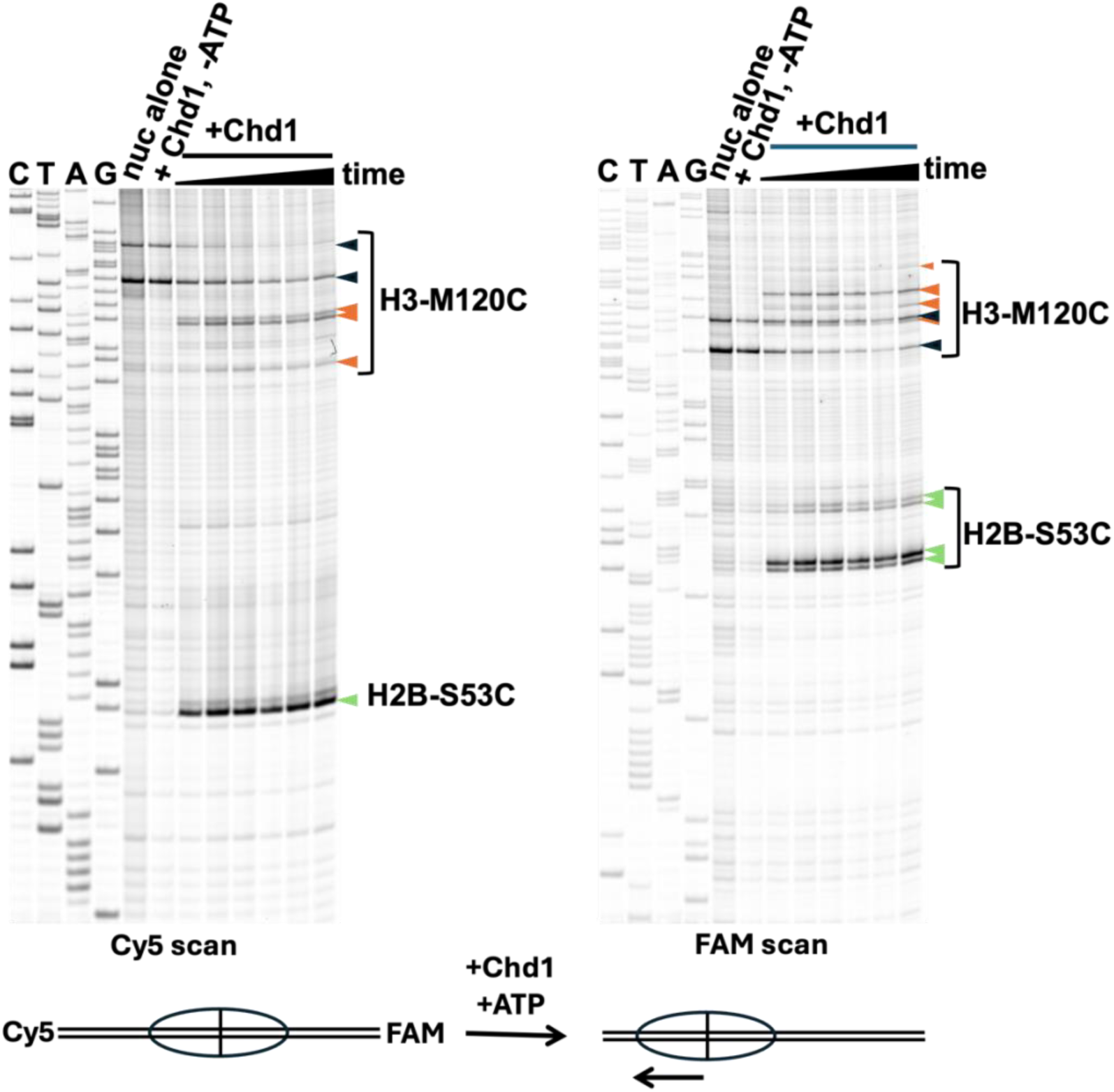
Mapping of 40-YNN1-40 during Chd1 remodeling. YNN1 nucleosomes (Cy5-40-YNN1-40-FAM) presenting cysteine residues on H2B-S53C and H3-M120C were mapped using APB crosslinking. Samples shown include nucleosome alone (black arrowheads) or with Chd1 (in a 1:1 stoichiometry) plus or minus ATP (orange and green arrowheads represent H3-M120C and H2B-S53C crosslinked sites, respectively, corresponding to shifted species). As monitored by the H3-M120C crosslinks, Chd1 showed a preference to reposition the octamer towards the 5’ side of the YNN1 dyad (in reference to the top strand). H2B-S53C did not show an APB crosslink in the nucleosome alone sample, likely due to the presence of an AT-rich sequence in the expected crosslinking region. Samples were visualized on a urea-denaturing gel and scanned for FAM and Cy5 on a Typhoon 5 imager.

### Supplemental Tables

**Table S1.**
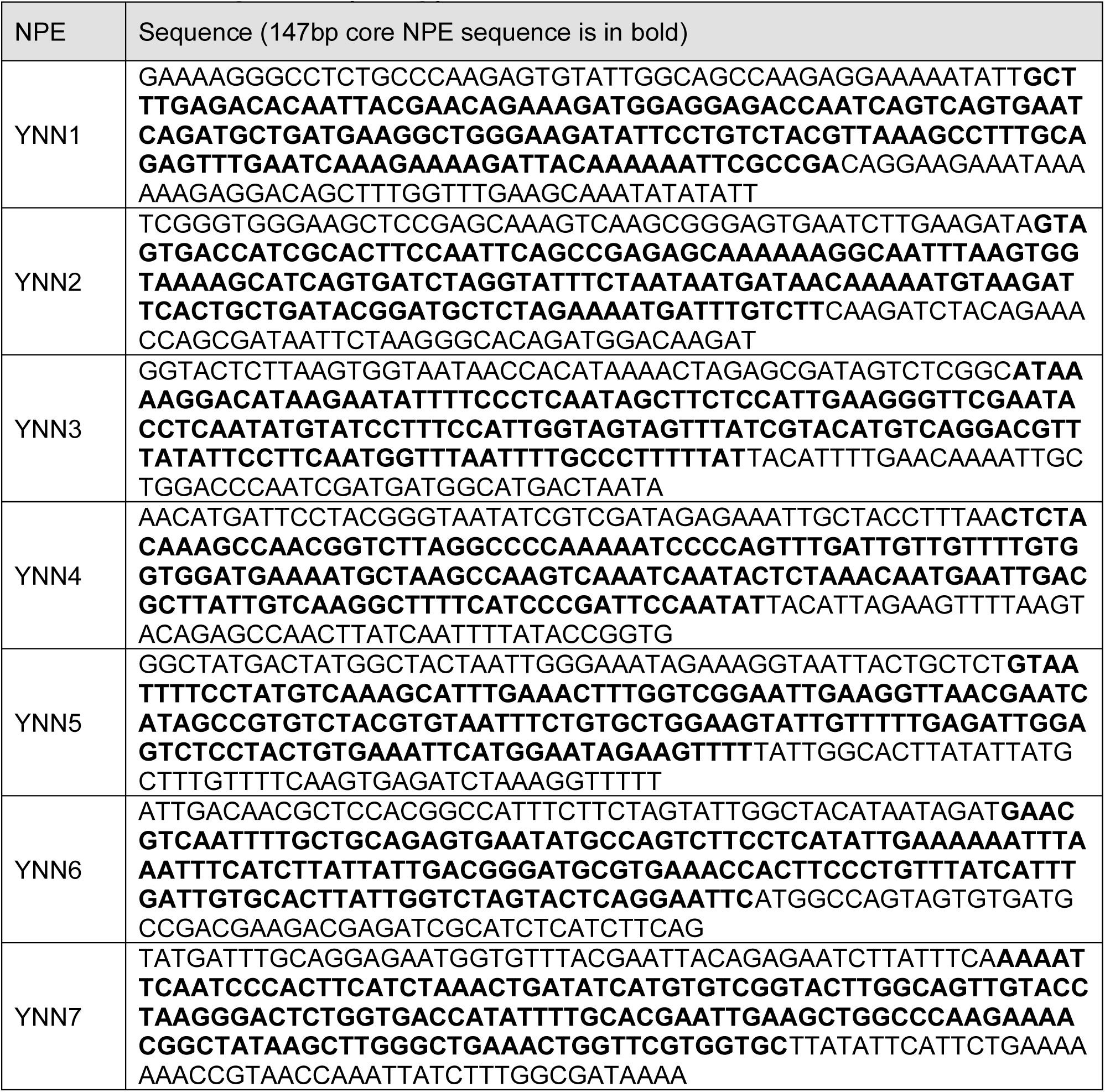
NPE sequences (247bp)

**Table S2.** Selected NPE candidates.

|  | chr | dyad in vivo | corr. in vivo | dyad in vitro | corr. in vitro |
| --- | --- | --- | --- | --- | --- |
| YNN1 | 16 | 349748 | 0.73182147 | 349743 | 0.92866253 |
| YNN2 | 10 | 219874 | 0.86830163 | 219874 | 0.95881774 |
| YNN3 | 4 | 676760 | 0.74084017 | 676765 | 0.86210489 |
| YNN4 | 2 | 276528 | 0.93791029 | 276548 | 0.73645788 |
| YNN5 | 7 | 633589 | 0.91721962 | 633589 | 0.97044051 |
| YNN6 | 10 | 502239 | 0.90942274 | 502224 | 0.90372847 |
| YNN7 | 12 | 426103 | 0.90149354 | 426088 | 0.90178566 |

**Table S3.**
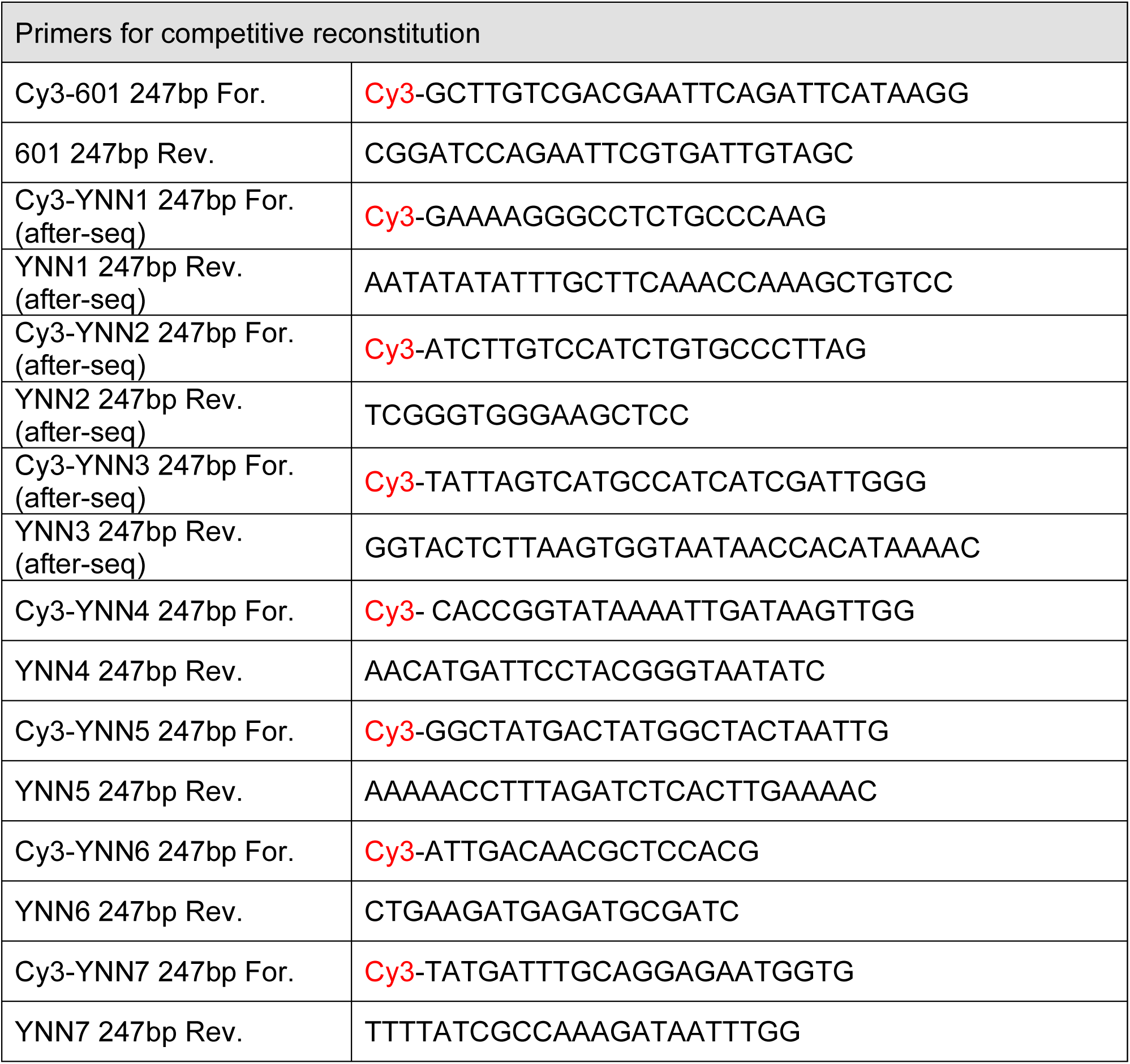
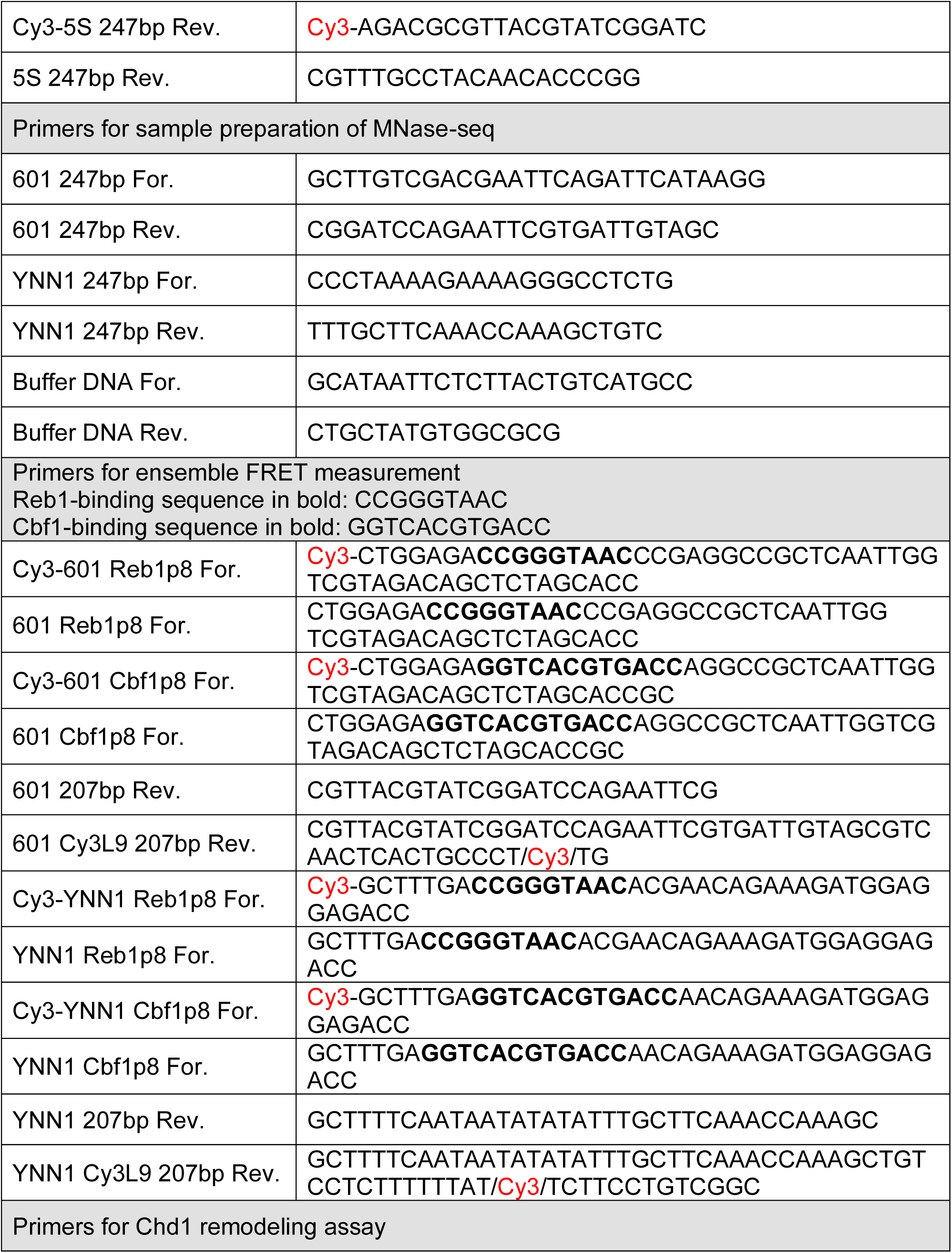
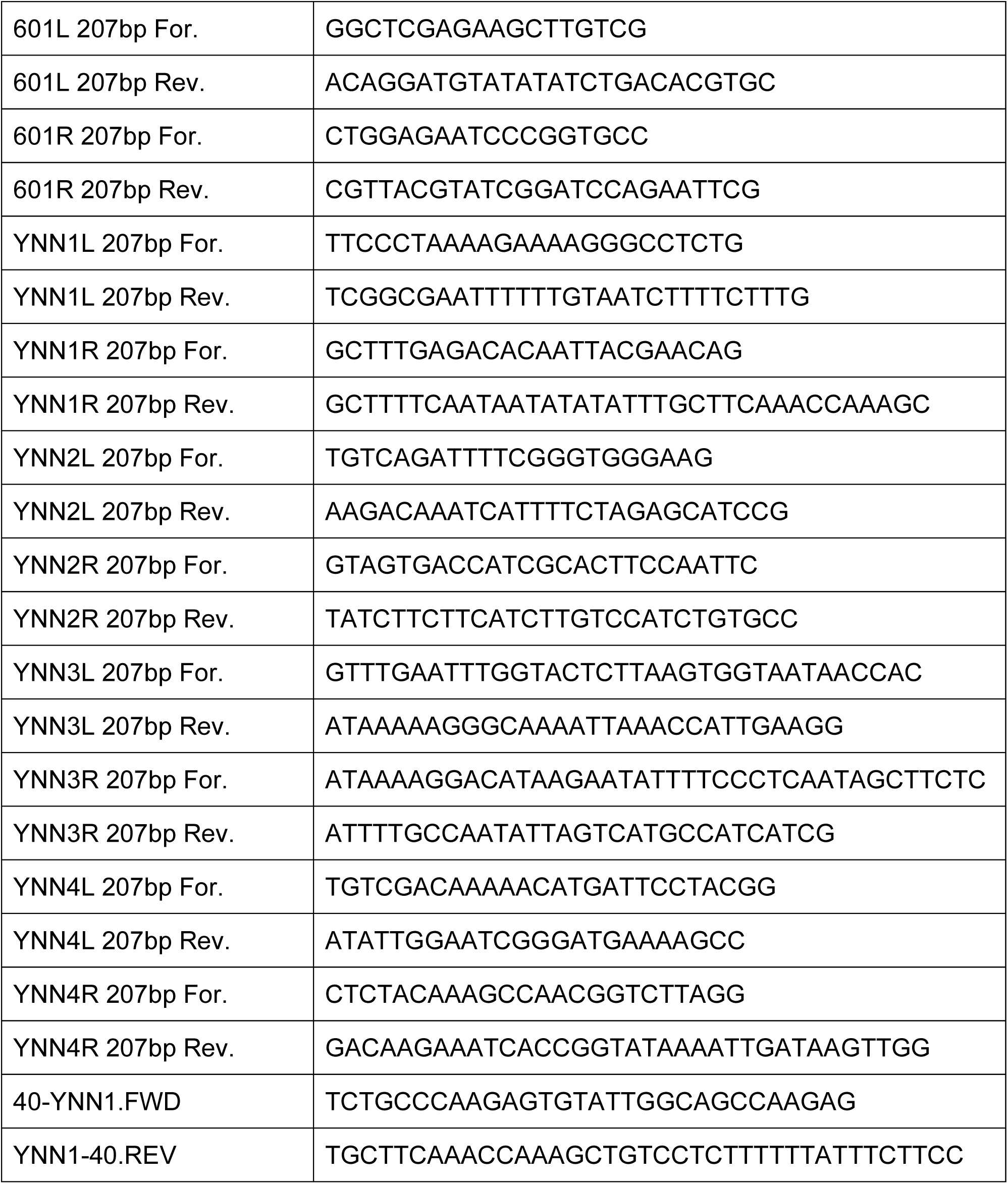
Primers used in the experiments.

**Table S4.** Gibbs free energy of DNA-histone interaction.

| | $\Delta\Delta G$ (kcal mol<sup>-1</sup>) |
| --- | --- |
| 5S DNA | 0 |
| YNN1 | 0.28 ± 0.07 |
| YNN2 | 0.48 ± 0.01 |
| YNN3 | 0.16 ± 0.09 |
| YNN4 | 0.3 ± 0.1 |
| YNN5 | 0.28 ± 0.08 |
| YNN6 | 0.37 ± 0.05 |
| YNN7 | 0.40 ± 0.04 |
| Widom 601 | -1.78 ± 0.05 |

**Table S5.** Distribution of DNA length population after MNase digestion.

| 601 (Reb1) |  |  |  |  |
| --- | --- | --- | --- | --- |
|  | 0 nM |  | 60 nM |  |
|  | replicate 1 | replicate 2 | replicate 1 | replicate 2 |
| 120 bp - 140 bp | 5.6% | 19.6% | 4.4% | 20.2% |
| 140 bp - 160 bp | 32.5% | 58.3% | 31.1% | 60.5% |
| 160 bp - 180 bp | 55.5% | 15.5% | 54.7% | 13.8% |
| 180 bp – 200 bp | 6.5% | 1.1% | 7.5% | 1.1% |
| YNN1 (Reb1) |  |  |  |  |
|  | 0 nM |  | 60 nM |  |
|  | replicate 1 | replicate 2 | replicate 1 | replicate 2 |
| 120 bp - 140 bp | 36.0% | 38.0% | 24.2% | 26.7% |
| 140 bp - 160 bp | 38.9% | 37.6% | 46.4% | 53.9% |
| 160 bp - 180 bp | 3.7% | 3.0% | 12.1% | 5.0% |
| 180 bp – 200 bp | 0.5% | 0.2% | 2.6% | 2.5% |
| YNN1 (Cbf1) |  |  |  |  |
|  | 0 nM |  | 60 nM |  |
|  | replicate 1 | replicate 2 | replicate 1 | replicate 2 |
| 120 bp - 140 bp | 20.3% | 33.0% | 22.8% | 27.1% |
| 140 bp - 160 bp | 56.5% | 46.9% | 54.5% | 50.0% |
| 160 bp - 180 bp | 15.1% | 8.6% | 13.9% | 13.2% |
| 180 bp – 200 bp | 1.6% | 1.2% | 1.4% | 2.4% |
| YNN1 (Pho4) |  |  |  |  |
|  | 0 nM |  | 60 nM |  |
|  | replicate 1 | replicate 2 | replicate 1 | replicate 2 |
| 120 bp - 140 bp | 20.6% | 20.0% | 18.3% | 15.8% |
| 140 bp - 160 bp | 50.6% | 47.3% | 50.9% | 52.2% |
| 160 bp - 180 bp | 12.9% | 14.1% | 16.0% | 19.1% |
| 180 bp – 200 bp | 1.6% | 2.2% | 3.8% | 5.0% |

**Table S6.**
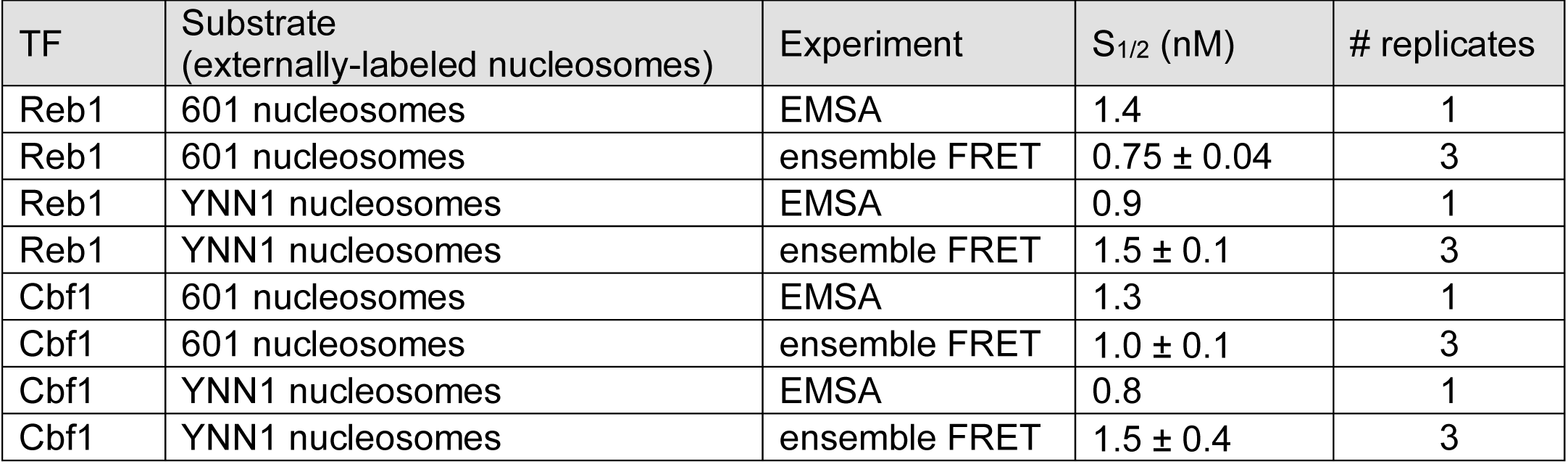
Binding affinities measured in the experiments.

**Table S7.** Chd1 remodeling rate in different YNN candidates.

|  | direction | rate (min<sup>-1</sup>) | P value |
| --- | --- | --- | --- |
| Widom 601 | fast | 0.31 ± 0.04 | 0.02 |
|  | slow | 0.08 ± 0.01 |  |
| YNN1 | fast | 0.26 ± 0.03 | 0.08 |
|  | slow | 0.16 ± 0.03 |  |
| YNN2 | fast | 0.23 ± 0.03 | 0.46 |
|  | slow | 0.19 ± 0.03 |  |
| YNN3 | fast | 0.33 ± 0.08 | 0.13 |
|  | slow | 0.08 ± 0.02 |  |
| YNN4 | fast | 0.33 ± 0.08 | 0.98 |
|  | slow | 0.27 ± 0.09 |  |

**Table S8.** Table of fractions remodeled by Chd1 in different YNN candidates.

|  | direction | fraction | P value |
| --- | --- | --- | --- |
| Widom 601 | Fast (right) | 0.60 ± 0.02 | 0.09 |
|  | Slow (left) | 0.53 ± 0.02 |  |
| YNN1 | Fast (left) | 0.77 ± 0.02 | 0.001 |
|  | Slow (right) | 0.37 ± 0.01 |  |
| YNN2 | Fast (left) | 0.59 ± 0.01 | 0.27 |
|  | Slow (right) | 0.54 ± 0.02 |  |
| YNN3 | Fast (right) | 0.54 ± 0.02 | 0.20 |
|  | Slow (left) | 0.47 ± 0.03 |  |
| YNN4 | Fast (left) | 0.53 ± 0.02 | 0.80 |
|  | Slow (right) | 0.54 ± 0.03 |  |

